# Positive Interpretation of Emotional Ambiguity Across Development: LC–dlPFC Circuitry

**DOI:** 10.64898/2026.06.10.730879

**Authors:** Tae-Ho Lee, Ya-Yun Chen, Joshua Neal

**Author notes:** Correspondence should be addressed to Tae-Ho Lee, Department of Psychology at Virginia Tech, 890 Drillfield Drive, Virginia Tech, Blacksburg, VA 24060.

## Abstract

Emotionally ambiguous facial expressions provide a tractable model for studying how uncertain affective information is resolved into categorical judgments. Here we tested whether developmental differences in this process are reflected in locus coeruleus (LC) circuitry and how these neural effects relate to behavioral and decision-process indices of interpretive bias. Participants from late childhood through middle-aged adulthood completed an angry-to-happy facial morph task during fMRI. Behavioral responses were analyzed using signal detection theory, and drift-diffusion modeling, and fMRI data were analyzed using LC-seeded generalized psychophysiological interaction and trial-level neural models. Positive bias increased systematically across development. Drift-diffusion analyses showed that this shift was captured specifically by drift intercept, a stimulus-independent evaluative component of evidence accumulation, rather than by perceptual sensitivity, starting point, or response caution. At the neural level, the association between LC–dlPFC coupling and positive bias was strongest in middle-aged adults. Trial-level analyses further indicated that dlPFC carried the clearer moment-to-moment evaluative signal, whereas LC contributed through slower temporal properties, indexed by the Hurst exponent, that shaped bias-related cortical coupling. Importantly, higher LC structural integrity (contrast-to-noise ratio) was associated with lower LC temporal persistence and, in turn, more negative LC–dlPFC coupling. Together, these findings suggest that age-related differences in positive interpretation are not simply differences in perceptual sensitivity or response tendency, but reflect age-graded variation in evaluative evidence accumulation supported by LC–dlPFC circuitry. This work extends recent evidence implicating LC–prefrontal connectivity in ambiguity processing and identifies evaluative drift bias as an age-graded circuit-level signature of emotional ambiguity resolution.

**Significance Statement:** How the brain resolves emotionally ambiguous faces across development is poorly understood. Using psychometric, drift-diffusion, and fMRI analyses in youth, emerging adults, and middle-aged adults, we show that age-related increases in positive interpretation are explained by drift intercept, a stimulus-independent evaluative component of evidence accumulation, rather than by perceptual sensitivity or starting-point bias. LC–dlPFC coupling tracked this same bias dimension. Trial-level analyses suggested that dlPFC carried the clearer moment-to-moment evaluative signal, whereas LC contributed through slower temporal properties of bias-related coupling. These findings identify evaluative drift bias as a measurable circuit-linked mechanism of emotional ambiguity resolution.

## Introduction

Perceiving emotion from a face is rarely a purely sensory act. Even when facial features are held constant, ambiguous emotional expressions can be categorized differently depending on observer characteristics and context (Ko et al., 2011; Lee et al., 2012; Lee et al., 2020). Age is one important dimension of this variability: the positivity effect describes a later-life shift in affective processing toward positive over negative information (Mather & Carstensen, 2005). A recent study linked this positive interpretive bias to the locus coeruleus (LC), showing that older adults exhibited greater LC activity during ambiguous-face resolution and stronger LC–dorsolateral prefrontal cortex (dlPFC) coupling, with both neural responses associated with more positive ambiguity resolution (Dave et al., 2025). This finding converges with recent work specifying how top-down influences, including attentional gain and predictive expectations, shape emotion perception (Mohanty et al., 2025). LC–dlPFC circuitry is well positioned to support this process: LC regulates cortical gain and arousal state through tonic and phasic modes (Aston-Jones & Cohen, 2005; Sara, 2009), whereas dlPFC supports controlled appraisal processing (Buhle et al., 2014; Ochsner & Gross, 2005).

However, because current evidence linking LC–dlPFC circuitry to positive ambiguity resolution comes primarily from older adults, it remains unclear whether this circuit reflects later-life compensatory recruitment (e.g.,Reuter-Lorenz & Cappell, 2008) or a broader mechanism in which LC-linked arousal signals interact with prefrontal evaluative processes to bias interpretation under uncertainty. These accounts make different developmental predictions: a compensation account predicts that the coupling–bias association should emerge mainly when age-related prefrontal inefficiency is prominent, whereas a general-mechanism account predicts that this association should be observable before late life. Testing participants from childhood through midlife therefore provides a way to separate these accounts, because LC and prefrontal systems both change across the lifespan (Aboud et al., 2019; Dave et al., 2026; Jacobs et al., 2018; Mather, 2020; Song et al., 2021).

The present study asked whether positive interpretation of emotional ambiguity prior to older adulthood is linked to LC–dlPFC circuitry. A developmental shift toward positive interpretation can in principle arise from several distinct sources: sharper perceptual discrimination, a pre-evidence response bias, a stimulus-independent evaluative bias in evidence accumulation, a change in response caution, or a change in non-decision processes(Mohanty et al., 2025). Without separating these components, it is not possible to ask which decision process LC–dlPFC circuitry tracks. To this end, we estimated individual differences in interpretive bias and related decision-process components using signal detection theory (Macmillan, 2002) and the drift-diffusion model (Ratcliff, 1978; Ratcliff & McKoon, 2008), and tested how each component related to LC–dlPFC connectivity. The sample included participants approximately 8 to 54 years of age, allowing us to test whether LC–dlPFC coupling is expressed before older adulthood and whether its association with positive interpretive bias differs across developmental groups. To further characterize LC contributions, we also examined neuromelanin-sensitive LC contrast as a putative proxy for LC structural integrity and LC signal persistence indexed by the Hurst exponent. Finally, we examined emotion regulation as a behavioral correlate of this process, because cognitive reappraisal recruits prefrontal control systems, including dlPFC, to reshape affective interpretations under uncertainty (Buhle et al., 2014; Gross & John, 2003; Ochsner et al., 2012). If LC–dlPFC circuitry supports top-down ambiguity resolution, reappraisal should relate to the same decision component and circuit-level signal associated with positive interpretive bias. Across these analyses, we find that developmental increases in positive interpretation are captured specifically by the stimulus-independent component of evidence accumulation, that LC–dlPFC coupling tracks this same evaluative-bias dimension, and that LC structural and temporal signal properties relate to the bias-defined coupling phenotype. Together, these findings extend LC–prefrontal ambiguity-resolution circuitry beyond older adulthood and identify evaluative drift bias as a circuit-linked target of emotional ambiguity resolution.

## Methods

### Participants

A total of 95 participants completed the forced-choice face perception task and were considered for the present analyses. Participants were divided into three developmental groups: children/adolescents (CH: N=34, ages 8–17 years, M=12.0 ± 2.5; 47.1% female), emerging adults (EA: N=28, ages 18–26 years, M=20.7 ± 1.5; 50.0% female), and middle-aged adults (MA: N=33, ages 33–54 years, M=42.7 ± 5.9; 57.6% female). All participants provided written informed consent, or assent with parental consent in the case of minors, and the study was approved by the Institutional Review Board. Five participants were excluded for behavioral data quality: one MA with an extrapolated PSE outside the presented morph range, three CH with unstable logistic fits (R² < .80), and one CH with a truncated task run (<50% planned trials). This yielded a final analyzable behavioral sample of 90 participants (CH 31; EA 28, MA 31; overall age range=8–54 years, 52.2% female).

### Task and Stimuli

Participants completed a forced-choice emotion-categorization task using faces morphed along an angry-to-happy continuum. Each trial began with a 0.5s fixation cross, followed by a face presented for up to 2.7s or until response. Intertrial intervals were jittered between 1-3.5s (**Fig 1A**). On each trial, participants judged whether a face appeared positive or negative. Stimuli consisted of 14 identities, 7 female and 7 male, drawn from established facial-expression datasets (Lundqvist et al., 1998; Tottenham et al., 2009). Angry and happy expressions were morphed with neutral expressions, and three intensities per emotion were selected (15%, 35%, 60%) plus neutral. This yielded seven morph levels: AN60, AN35, AN15, NE, HA15, HA35, and HA60 (**Fig S1**). Each of the 98 unique stimuli was presented twice.

**Fig. 1.**
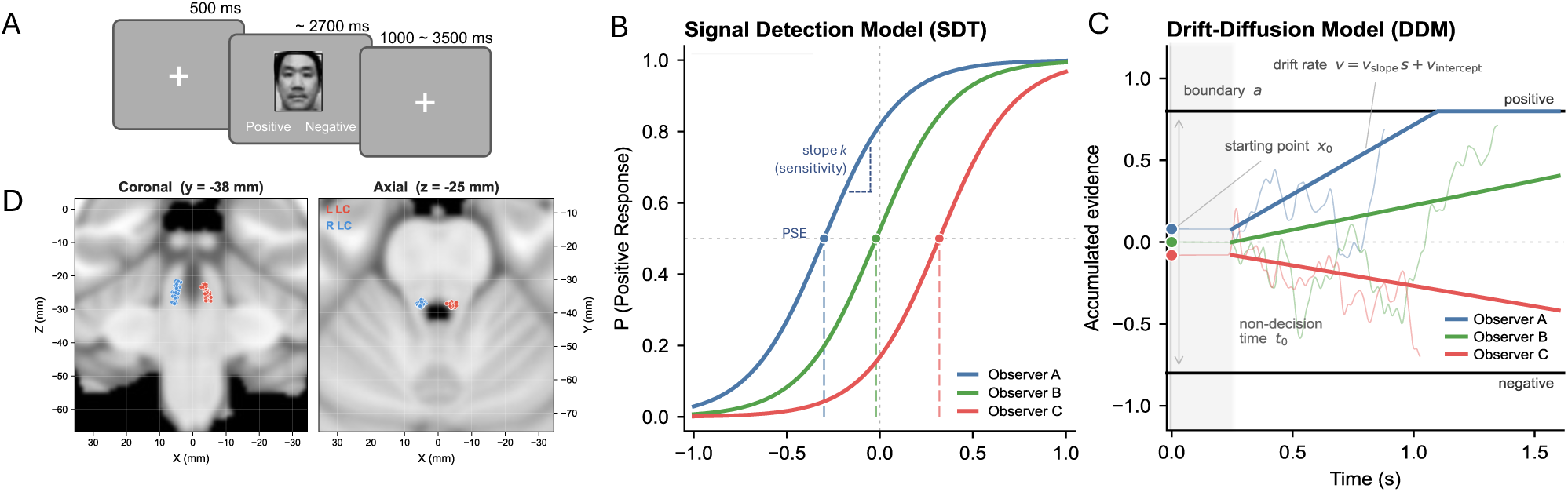
Schematic of psychometric and drift-diffusion accounts of positive ambiguity resolution, with LC localization. (A) Schematic of the forced-choice emotion categorization task using morphed facial expressions spanning angry to happy intensity levels (. Example faces shown are illustrative and were included only to convey the task structure; they were not the actual stimuli used in the experiment. (B) Signal detection model (SDT). Psychometric functions for three hypothetical observers differing in positive interpretive bias. Observer A (blue) has a left-shifted category boundary, indicating stronger positive bias; Observer B (green) is approximately unbiased; and Observer C (red) has a right-shifted boundary, requiring more clearly happy stimuli before responding “positive.” The slope parameter k is held constant across observers for illustration and indexes perceptual sensitivity.(C) Drift-diffusion model (DDM). Simulated evidence-accumulation trajectories for the same three hypothetical observers at a fixed, mildly ambiguous stimulus. Observer A accumulates evidence toward the positive decision bound, *B*, reflecting a more positive drift rate and a slightly positive starting point (x₀ > 0), whereas Observer C accumulates toward the negative bound, *−B*. Drift rate is parameterized as v = v*_slope_* × intensity + v_intercept_, where v*_slope_* indexes perceptual gain and v_intercept_ indexes stimulus-independent evaluative drift bias. Boundary separation, *B*, sets the amount of evidence required before committing to a response, and non-decision time (t; shaded gray) captures stimulus encoding and motor execution. (D) **LC localization.** Individual left and right LC peak voxels identified from neuromelanin-sensitive imaging are overlayed in representative coronal and axial brainstem views and projected onto the MNI152 template for the full sample (blue = right LC, red = left LC). Note. Photographs shown in the task schematic are of the authors and are reproduced with their permission.

### Questionnaires

All participants completed the Emotion Regulation Questionnaire (ERQ; Gross & John, 2003), which assesses habitual use of cognitive reappraisal (6 items) and suppression (4 items). We computed an ER composite score as z(reappraisal) − z(suppression), with higher scores indicating greater relative reliance on reappraisal than suppression. We used the composite as the primary ER measure to retain a single index across analyses; subscale-level (reappraisal, suppression) results are reported in Supplementary Fig S4P–T for completenes

### Signal Detection Theory Model

#### Perceptual Threshold and Sensitivity Estimation

For each participant, the proportion of “positive” responses at each morph level was fitted with a two-parameter logistic psychometric function, P(x)=1 / (1 + exp(−k(x − PSE))) (**Fig 1B**). The point of subjective equality (PSE) indexed the morph level at which positive and negative responses were equally likely. Because higher PSE values indicate that more overt happiness was required for a positive judgment, positive bias was defined as 1 − PSE, such that higher values indicate a lower threshold for positive categorization. The slope parameter *k* indexed perceptual sensitivity, with larger values indicating a sharper discrimination sensitivity between intensities.

To stabilize the endpoint behavior of the logistic fit and reduce unstable solutions near the floor and ceiling (Prins, 2019), we appended two theoretical anchor points: 100% angry (P=0) and 100% happy (P=1). These anchors corresponded to fully unambiguous expressions expected to lie at the floor and ceiling of the response function, respectively (cf. Wang et al., 2017). Logistic functions were fitted for each participant to nine points along the morph continuum using nonlinear least-squares optimization (PSE ∈ [0, 1], k ∈ [0.1, 50.0]). As a sensitivity check, we also fitted the same function to the seven empirical morph levels alone. Seven-level-only fits produced nearly identical estimates (PSE: r=.997; k: r=.979), confirming that anchor stabilization did not alter individual-difference estimates; 9-point fits were therefore used in primary analyses.

#### Subject-Specific Maximum-Ambiguity Level

To account for individual differences in perceived emotional ambiguity, we defined each participant’s maximum-ambiguity condition from their fitted PSE (**Fig S1**). The continuous PSE value was transformed to the seven-level morph scale using the formula scale=(PSE − 0.5) × 6 and assigned to the nearest discrete morph condition. This PSE-nearest condition was treated as the participant’s point of maximum subjective ambiguity and was used for participant-specific gPPI and task-evoked GLM contrasts, rather than relying on a fixed stimulus-defined ambiguity midpoint. In the final sample, PSE-nearest morph conditions were distributed as follows: HA15 (57.8%), HA35 (16.7%), NE (11.1%), AN15 (10.0%), and AN35 (4.4%).

### Drift-Diffusion Model

To complement the psychometric characterization, we fitted a drift-diffusion model (DDM; Fig. 1C; Ratcliff, 1978; Ratcliff & McKoon, 2008) to each participant’s trial-level reaction time and choice data using pyDDM (Shinn et al., 2020). Trial-by-trial drift rate was parameterized as v=v_slope_ × s + v_intercept_, where s denotes signed morph intensity, rescaled to [−0.60, 0.60] with neutral=0. Drift slope (v_slope_) captures how strongly evidence accumulation changes with stimulus intensity and therefore indexes perceptual sensitivity or gain, corresponding conceptually to the psychometric slope *k*. Drift intercept (v_intercept_) indexed stimulus-independent accumulation toward the positive boundary and was treated as the DDM counterpart of psychometric positive bias: both quantify whether ambiguous faces are resolved more positively or negatively. Boundary separation (B) reflects response caution, with larger values indicating that more evidence is required before a decision. Starting point (x₀) captures an initial bias before evidence accumulation begins, indicating whether the decision process starts closer to one response boundary. Non-decision time (t) captures processes outside the decision itself, including stimulus encoding and motor execution. Noise was fixed at σ=1, and a small uniform mixture component was included in the response-time density to absorb contaminant trials. Trials with reaction times <0.2s or >2.7s, as well as trials with no recorded response, were excluded before fitting. Five free parameters were estimated by maximum-likelihood optimization within the following bounds: v_slope_ ∈ [0.5, 30], v_intercept_ ∈ [−3, 3], B ∈ [0.3, 4.0], x0 ∈ [−0.7, 0.7], and t ∈ [0.1, 0.6] s.

### fMRI Acquisition and Preprocessing

All MRI data were acquired on a Siemens 3T PRISMA with a 64-channel matrix head coil. High-resolution T1w (TR=2.5 s; TE=2.06 ms; FA=8°; 1 mm isotropic voxel) and T2w (TR=3.2 s; TE=563 ms; FA=120°; 1 mm isotropic voxel) anatomic images were acquired for tissue segmentation (GM, WM, and CSF mask) and normalization. Functional images were acquired with gradient-echo echo-planar T2*-weighted imaging sequence (TR=2s; TE=25 ms; FA=90°; 2.5 x 2.5 mm resolution; 37 interleaved 3.0 mm slices with 0.3 mm gap). Neuromelanin-sensitive MRI was collected using a T1-weighted Fast Spin Echo imaging sequence (repetition time=750 ms, echo time=12 ms, flip angle=120°, 11 axial slices, bandwidth=285Hz/Px, slice thickness=2.5 mm, in-plane resolution=0.43×0.43 mm). Preprocessing was performed using the FMRIB Software Library (FSL; Jenkinson et al., 2012), ICA-AROMA toolbox (Pruim et al., 2015), and ANTs library (Avants et al., 2009). ICA-AROMA was used to reduce motion-related and physiological noise in the LC signal when simultaneous physiological recordings were unavailable (Prokopiou et al., 2022; Scheel et al., 2022). Preprocessing included the first two volumes cut, high pass filter (128s;), motion correction, 5-mm smoothing, grand-mean intensity normalization, ICA denoising, and registration to standard MNI 2 mm brain template.

### LC-Seed Definition and Structural Integrity Calculation Based on Neuromelanin Imaging

For each participant, the LC seed used in the gPPI analysis was defined from the neuromelanin-sensitive image as the highest-intensity voxel within the LC (**Fig 1D**). Because LC neuromelanin signal is strongest near the core, the peak-intensity voxel was used as a subject-specific LC estimate. Each neuromelanin image was registered to the 1-mm neuromelanin MNI template, and the corresponding probabilistic LC atlas (Lee et al., 2024) was inverse-warped to native space. Within each hemisphere, the LC mask was thresholded at 50% probability, the peak-intensity voxel was selected. This procedure has been validated against manual LC tracing in prior work (Neal et al., 2025). Finally, the LC seed was defined as 2-mm sphere on this peak. Sensitivity analyses repeated the whole-brain gPPI using 3-mm and 4-mm peak-centered spheres and the 50% probabilistic LC atlas seed (**Fig S2**).

Using each participant’s LC peak voxel, we computed a contrast-to-noise ratio (CNR) for each participant as (LC peak − PT mean) / PT mean, where PT denotes the pontine tegmentum reference region (Neal et al., 2025; Sasaki et al., 2006; Shibata et al., 2006). Neuromelanin-sensitive images were unavailable for two CH and one MA because of scan-time constraints; for these participants, the LC seed was defined from the 50% probabilistic atlas. CNR was therefore available for 87 of the 90 participants.

### LC-Seed Functional Connectivity Analysis

To examine task-dependent functional connectivity between the LC and cortical regions, we conducted a generalized psychophysiological interaction (gPPI) analysis. Left and right LC seeds were analyzed separately to preserve hemisphere-specific coupling profiles and avoid spatial mixing across the small bilateral nuclei (Chandler et al., 2014; Schwarz et al., 2015). For each hemisphere, seed time series were extracted from each participant’s preprocessed-but-unsmoothed native EPI data to preserve LC-specific signal and minimize spatial mixing, which can bias connectivity estimates in this small midbrain nucleus (Alakörkkö et al., 2017).

The individual-level GLM included seven event-related task regressors corresponding to the seven morph conditions. Each regressor was convolved with a double-gamma hemodynamic response function and paired with its temporal derivative. The gPPI model used the same first-level GLM design, with the LC seed time series added as the physiological regressor. For each morph condition, the psychophysiological interaction term was constructed as the element-wise product of the LC seed time series and the corresponding mean-centered task regressor, indexing condition-dependent changes in LC–cortical coupling.

For group-level inference, we used each participant’s gPPI contrast corresponding to their subject-specific maximum-ambiguity condition, defined as the morph condition nearest to that participant’s PSE. This ensured that the group-level analysis tested LC–cortical coupling during each participant’s point of maximal subjective ambiguity rather than during a fixed stimulus-defined condition. Group-level inference was performed using FSL FLAME1+2, with age included as a nuisance covariate and the behavioral variable of interest entered as the primary covariate. SDT-derived positive bias (1 − PSE) was used as the primary behavioral covariate. Parallel models using DDM drift intercept and other decision parameters are reported in Supplementary Materials (**Fig S3** and **Table S1**). Whole-brain statistical maps were thresholded at Z > 2.57 and cluster-corrected at p=.05. Significant group-level clusters were then used as ROI masks for follow-up analyses and developmental group comparisons.

### Trial-Level Drift-Diffusion Modeling with Regional BOLD Regressors

To estimate trial-level neural contributions to evidence accumulation, we conducted a drift-diffusion analysis in which right LC and left dlPFC activity were entered simultaneously as predictors of drift rate. To this end, we first estimated single-trial BOLD activity for each region using least-squares single-trial estimation (LSS ; Mumford et al., 2012). Within each participant, single-trial BOLD values for each region were winsorized at the 5th and 95th percentiles to attenuate the influence of extreme trial-level estimates, which can arise in single-trial estimation due to collinearity between the target-trial regressor and temporally adjacent trials, and then z-scored to remove between-subject differences in overall BOLD signal amplitude.

We then refit the model for each participant, retaining the same five baseline parameters (v*_slope_*, v*_intercept_*, B, x0, and t, plus a uniform mixture component) while adding three neural contributors to drift rate: one term for standardized single-trial dlPFC BOLD (β_D_) one term for standardized single-trial LC BOLD (β_L_), and one term for their product, indexing the trial-level LC × dlPFC interaction (β_I_). To allow each regional contribution to vary as a function of how close a given trial’s stimulus was to the participant’s own perceptual boundary, each of the three neural terms was additionally multiplied by a subject-specific ambiguity weight that peaked at each participant’s PSE and decayed smoothly with distance from it (Gaussian weighting, σ=1 unit in the ordinal cope-selection scale introduced earlier). Formally, amb(t)=exp(−d(t)² / 2), where d(t) is the ordinal distance, in first-level contrast units, between the morph condition of trial t and the participant’s assigned PSE level. As a sensitivity analysis, we refit the same model using a linear ambiguity weighting term (1 − |distance|/3).

The three fitted coefficients (β_D_, β_L,_ β_I_) quantified baseline trial-level contributions of dlPFC activity, LC activity, and their interaction, whereas the three ambiguity-moderated coefficients (β_D×amb_, β_L×amb_, β_I×amb_) quantified whether each contribution changed as trials approached the participant’s perceptual boundary. All neural terms were entered into drift rate, and the baseline DDM parameters were re-estimated jointly with the neural terms using the same bounds as in the original fit.

### Temporal Persistence of LC Activity

The trial-level DDM analysis tested whether LC and dlPFC carry phasic, trial-by-trial contributions to drift rate. However, LC function is often characterized as setting tonic noradrenergic gain rather than modulating individual trials (Aston-Jones & Cohen, 2005). To test whether the LC signal operated at a sustained rather than trial-locked timescale, we computed the Hurst exponent of each participant’s LC seed time series using rescaled range (R/S) analysis (Hurst, 1951; You et al., 2012). The Hurst exponent indexes long-range temporal persistence in a signal. Values above 0.5 indicate temporally persistent dynamics, whereas values near 0.5 indicate uncorrelated fluctuations (Dong et al., 2018; Garrett et al., 2013; Mohr & Nagel, 2010). In the present context, and consistent with adaptive-gain accounts of LC function, higher Hurst values were interpreted as reflecting a more temporally persistent, tonic-like LC operating mode.

Because task state can modulate scale-free properties of the BOLD signal, we performed a sensitivity check by recomputing the Hurst exponent after regressing all seven HRF-convolved task-condition regressors from the LC time series. Task-residualized and full time-series estimates were nearly identical (r=.951), so full right LC Hurst values were used.

### Statistical Analysis

Unless otherwise noted, inferential statistics were based on 5,000 resamples. Group comparisons used permutation-based F tests followed by bootstrap pairwise mean-difference tests with Cohen’s d. Continuous associations were tested using bootstrap Pearson correlations and bootstrap partial correlations controlling for age. All tests were two-tailed and were conducted in Python using SciPy, statsmodels, and pyDDM.

## Results

### Signal Detection Theory Model Results

Group-mean psychometric functions across the seven morph levels are shown in **Fig 2A**. Positive bias increased systematically across developmental groups (F(2,87)=19.89, p<.001; **Fig 2B**). Middle-aged adults (M=0.494) showed greater positive bias than both children (M=0.293; 95% CI [0.139, 0.264], p<.001) and emerging adults (M=0.356; 95% CI [0.064, 0.209], p<.001). Children and emerging adults also differed reliably (95% CI [−0.115, −0.013], p=.010). The slope parameter, k, showed only a small group-level effect that did not reach conventional significance (F(2,87)=2.92, p=.058; **Fig 2C**). Thus, the main developmental signal in ambiguity resolution was expressed as a shift in positive interpretive bias rather than sharper perceptual discrimination. If anything, sensitivity was slightly reduced in the older group. Full-sample age associations are shown in **Fig S4A-B**.

**Fig 2.**
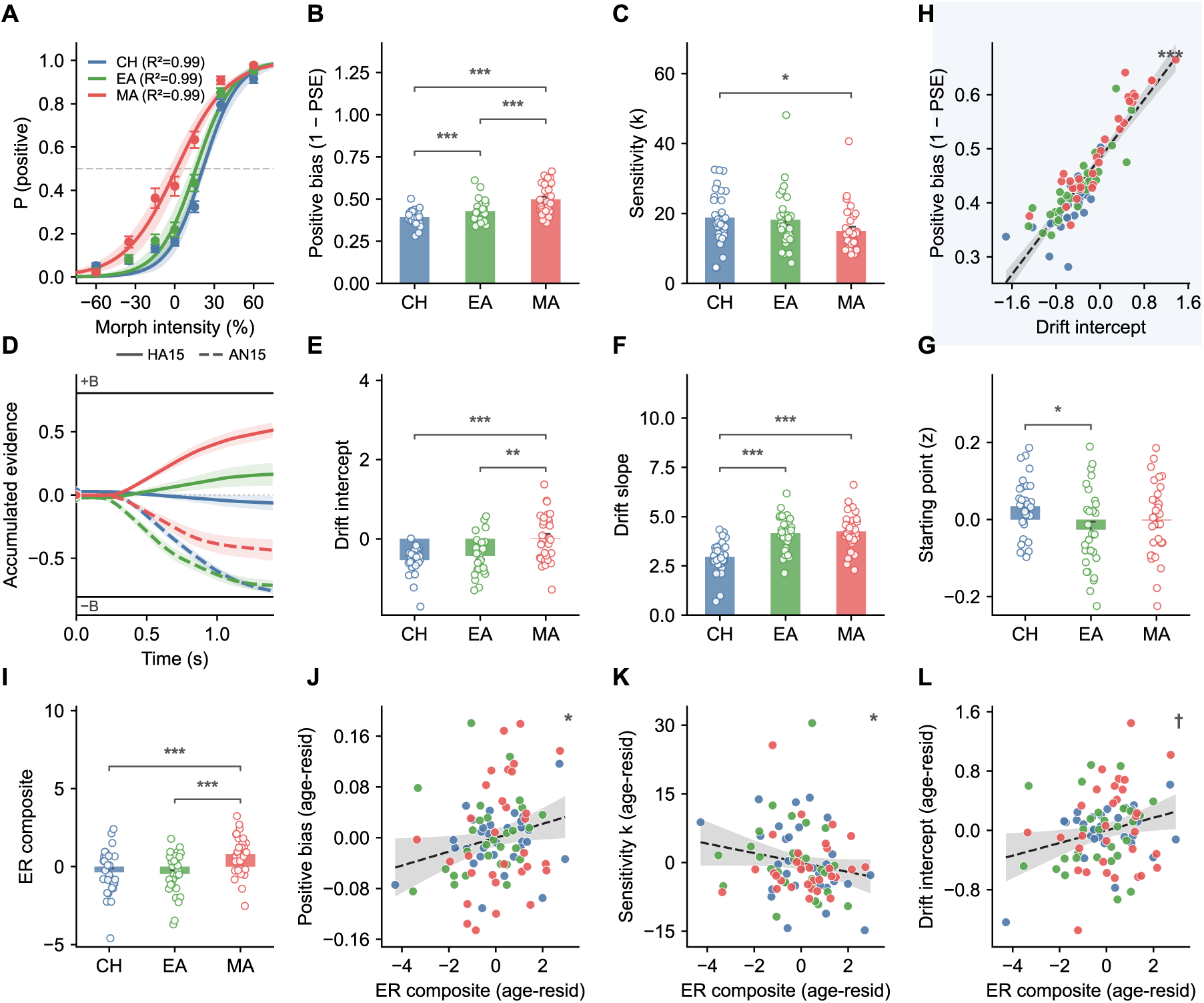
Signal detection, drift-diffusion, and emotion regulation characterization of ambiguity resolution across development. **Row 1: Signal Detection Model.** (A) Group-mean psychometric functions across seven morph intensities (AN60–HA60), with bootstrap 95% confidence bands. (B) Positive bias, indexing the evaluative shift in the categorization boundary, computed as 1 − PSE, by developmental group. (C) Sensitivity, indexing perceptual discriminability (slope *k* of the psychometric function), by group; middle-aged adults showed lower sensitivity than children (*p* = .011). **Row 2: Drift-Diffusion Model.** (D) Model-implied mean drift trajectories for each group, reconstructed from per-subject model fits (all seven stimulus intensities were fit jointly) and visualized here at two representative low-ambiguity levels: HA15 (15%; solid lines) and AN15 (−15%; dashed lines); shading around each line shows the SEM across subjects. (E) Drift intercept, indexing evaluative drift bias at a neutral stimulus, which is the DDM counterpart of 1 − PSE, by group. (F) Drift slope, indexing perceptual gain, defined as the per-unit increase in evidence accumulation with stimulus intensity, by group. (G) Starting point (*x*₀), indexing a pre-evidence response bias toward one choice boundary, by group **Row 3: Emotion Regulation and Its Behavioral Signatures.** (I) ER composite (z[reappraisal] − z[suppression]) by group. After controlling for age, the ER composite was associated with (J) higher positive bias, (K) lower perceptual sensitivity *k*, and (L) higher drift intercept, establishing a content-versus-gain dissociation across the two psychometric components of SDT and their direct DDM analogues. (H) Individual-level correspondence between SDT-derived positive bias and DDM drift intercept. The two measures were nearly identical (r = .908), indicating that the SDT and DDM frameworks captured the same evaluative-bias construct. Bars show group means ± SEM with individual data points overlaid. All scatter panels show age-residualized values. ^†^p = .052 \**p < .05, **p < .01, ***p < .001*.

### Drift-Diffusion Model Results

The DDM decomposed each participant’s choices and reaction times into five distinct components and revealed clear developmental dissociations between the perceptual-gain and evaluative components of evidence accumulation; group-mean model-implied drift trajectories are shown in **Fig 2D**. Drift intercept (v*_intercept_*), the stimulus-independent component of drift, closely tracked the developmental rise in positive bias and showed a significant group effect (F(2,87)=11.22, p<.001; **Fig 2E**). Middle-aged adults (M=0.015) showed a more positive drift intercept than both children (M=−0.533, 95% CI [0.314,0.780], p<.001) and emerging adults (M=−0.426; 95% CI [0.171,0.710], p=.002), with no reliable difference between children and emerging adults (p=.32). Thus, the evaluative component of the decision process continued to shift in a positive direction through middle adulthood.

Drift slope (v*_slope_*), indexing the per-unit increase in evidence accumulation with stimulus intensity, showed the largest group effect (F(2,87)=20.99, p<.001; **Fig 2F**). Children (M=2.951) had a markedly shallower drift slope than both emerging adults (M=4.152; 95% CI [−1.613,−0.778], p<.001) and middle-aged adults (M=4.244; 95% CI [−1.727, −0.894], p<.001), whereas the two adult groups did not differ (p=.71). This pattern suggests that developmental increases in perceptual gain occur primarily between childhood and early adulthood and then plateau.

Starting point (x0), indexing a pre-evidence bias toward one choice boundary that, in the canonical DDM framework, reflects a prior expectation about the upcoming stimulus (Mohanty et al., 2025; Ratcliff & McKoon, 2008), showed a modest group effect (F(2,87)=3.24, p=.042) Thus, although starting point varied modestly across groups, it did not account for the developmental increase in positive interpretive bias.

Taken together, these findings indicate that the DDM separates a perceptual-gain component that matures between childhood and emerging adulthood from an evaluative component that continues to shift through middle adulthood. The remaining DDM parameters and full-sample age associations are shown in **Fig S4C-I.**

### SDT–DDM Convergence

Across participants, SDT-derived positive bias and DDM drift intercept were nearly identical (r=.908, 95% CI [.863,.941], p<1e−32; **Fig 2H**), indicating that both models captured the same evaluative-bias dimension. Positive bias did not covary with drift slope, starting point, boundary separation, or nondecision time (all ps>.31; **Fig S4J–M**). Thus, the positive-bias effect was not consistent with a starting-point or response-caution account, but was localized to stimulus-independent drift.

### Reaction Time

Reaction times differed across groups at every morph intensity (all ps<.02), and followed the expected inverted-U pattern, with responses slowing near the category boundary. Emerging adults consistently responded fastest, whereas children responded slowest (**Fig S4N**).

### Emotion Regulation and Evaluative Bias

The ER composite differed across groups (F(2,87)=8.26, p<.001; **Fig 2I**). Middle-aged adults (M=.791) reported a more reappraisal-dominant profile than both children (M=−.364; 95% CI [0.501,1.802], p=.001) and emerging adults (M=−.473; 95% CI [0.635,1.950], p<.001), whereas children and emerging adults did not differ (p=.74). Full-sample age associations are shown in **Fig S4O.**

At the SDT level, the ER composite showed opposing associations with the two psychometric components: it was positively associated with positive bias (partial r | age=.229, 95% CI [.018,.421], p=.030; **Fig 2J**) and negatively associated with perceptual sensitivity (partial r | age=−.187, 95% CI [−.379,−.002], p=.048; **Fig 2K**). Thus, a more reappraisal-dominant regulation style was associated with stronger positive interpretive bias and a shallower psychometric slope. This SDT pattern aligns with the DDM finding that positive bias was primarily captured by drift intercept rather than drift slope, suggesting that reappraisal-dominant regulation is more closely related to evaluative interpretation than to perceptual gain.

DDM analyses showed a weaker but directionally consistent pattern. Drift intercept showed the clearest age-controlled association with the ER composite, although this effect was marginal (partial r | age=.243, p=.052; **Fig 2L**). Drift slope, boundary separation, and nondecision time were not reliably associated with the composite, and starting point showed only a borderline association that did not track positive bias.

Subscale analyses are reported in **Fig S4P–T** and indicated that the SDT association was more apparent for cognitive reappraisal than for expressive suppression.

### *Whole-Brain gPPI Analysis:* LC Connectivity Tracks Positive Interpretive Bias

We tested whether task-modulated LC functional connectivity was associated with individual differences in positive interpretive bias. In the whole-brain gPPI analysis using the right LC seed, the participant-specific maximum-ambiguity contrast revealed a significant dorsal left prefrontal cluster when age was included as a nuisance covariate. This cluster was centered in the superior frontal gyrus and extended into superior dlPFC, consistent with BA9/46 (peak MNI: −22, 34, 48; 245 voxels). Greater positive bias was associated with lower right LC–left dlPFC connectivity estimates (**Fig 3A**). This pattern is broadly consistent with recent evidence implicating LC–dlPFC connectivity in ambiguity resolution (Dave et al., 2025). The left LC seed identified a separate right angular gyrus/inferior parietal cluster near the temporoparietal junction (peak MNI: 44, −60, 32; 232 voxels). Because the primary goal of the present study was to test the LC–dlPFC pathway in relation to emotional ambiguity resolution, this secondary parietal finding is reported in the Supplementary Materials rather than pursued as a second main circuit (**Fig S5** and **Table S2**).

**Fig 3.**
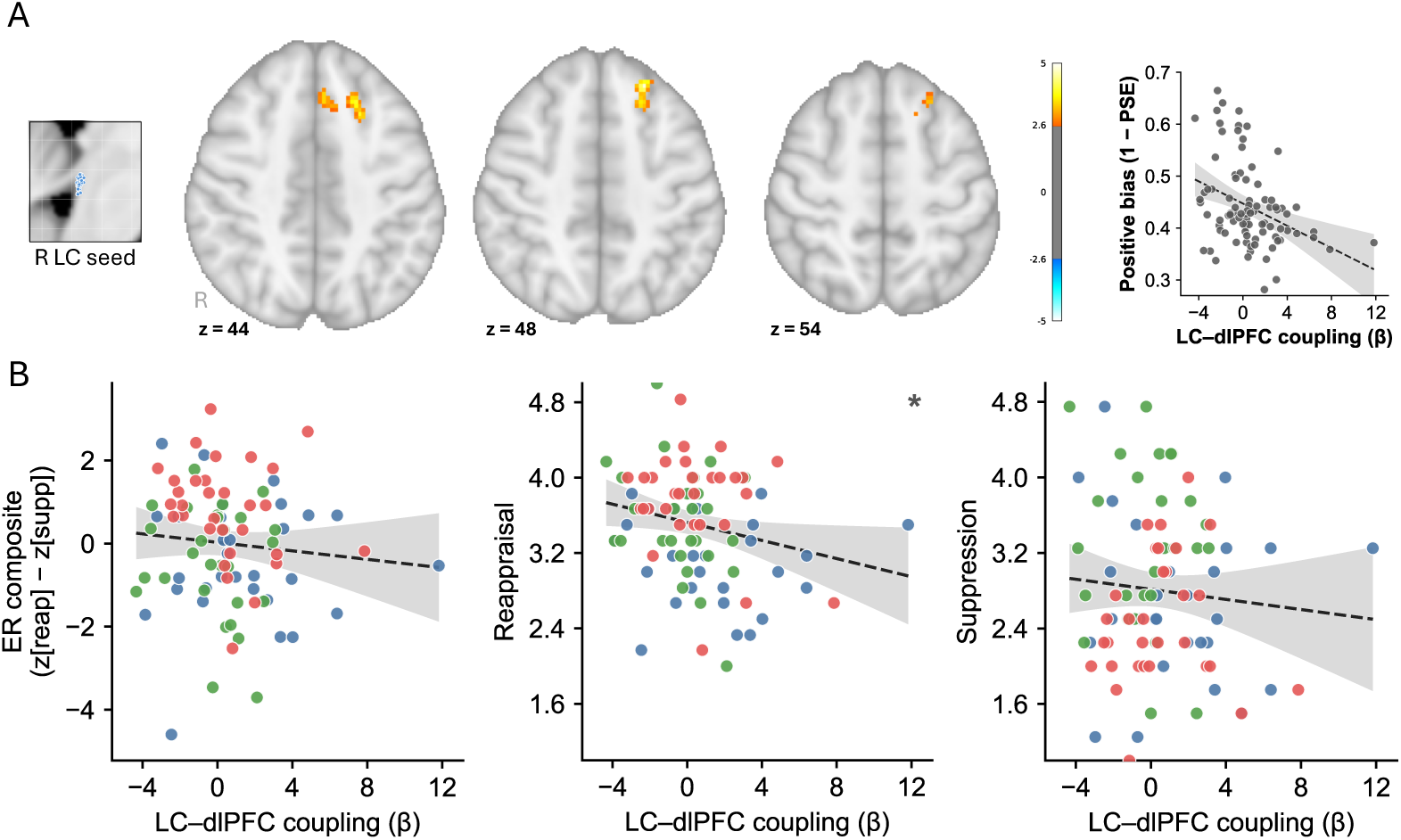
Right LC–Left dlPFC connectivity associated with positive bias and reappraisal. (A) Whole-brain gPPI analysis using the right LC seed and positive bias as the individual-difference covariate, with age included as a nuisance covariate. Axial slices show a significant cluster in the left superior/middle frontal cortex, consistent with superior dlPFC (peak = −22, 34, 48; 245 voxels; Z > 2.57, cluster-corrected p < .05). Scatterplot is shown for visualization of the extracted cluster effect (B) ROI analysis of extracted right LC–left dlPFC gPPI estimates plotted against emotion-regulation measures. Coupling was not reliably associated with the ER composite or expressive suppression, but was inversely associated with cognitive reappraisal. Color coding: CH = blue, EA = green, MA = red. ^†^p = .076, *p < .05, **p < .01, ***p < .001.

### *ROI Analysis:* LC–dlPFC Connectivity Tracks Reappraisal But Not Suppression

Within the same bias-defined dlPFC connectivity cluster, we tested whether emotion-regulation measures were reflected in LC–dlPFC connectivity (**Fig 3B**) The ER composite was not reliably associated with LC–dlPFC connectivity after controlling for age (p=.431). In subscale analyses, cognitive reappraisal showed an inverse association with LC–dlPFC connectivity (partial r | age=−.221, 95% CI [−.407,−.028], p=.024), whereas expressive suppression did not (p=.381). Thus, although the composite ER measure did not relate reliably to LC–dlPFC connectivity, the subscale analysis suggests that the bias-related LC–dlPFC circuit was more closely linked to cognitive reappraisal than to suppression.

### LC versus dlPFC as trial-level neural correlates of ambiguity resolution

The whole-brain gPPI analysis identified LC–dlPFC coupling as a bias-related circuit at the between-person level, but gPPI indexes task-related covariation between regions and does not specify whether trial-level activity in either region is more closely associated with evidence accumulation on individual decisions. To examine this, we refit the DDM for each participant with both regions entered simultaneously as trial-level regressors on drift rate, using LSS-derived single-trial BOLD estimates from each participant’s right LC seed and from the left dlPFC cluster identified in the primary gPPI analysis. Each region’s contribution was additionally modulated by a trial-level ambiguity weight centered on the participant’s PSE, yielding six per-subject β parameters: a main effect and an ambiguity-moderated term for LC (β_L_, β_L·amb_), dlPFC (β_D_, β_D·amb_), and their interaction (β_I_, β_I·amb_; **Fig 4A**).

**Fig. 4.**
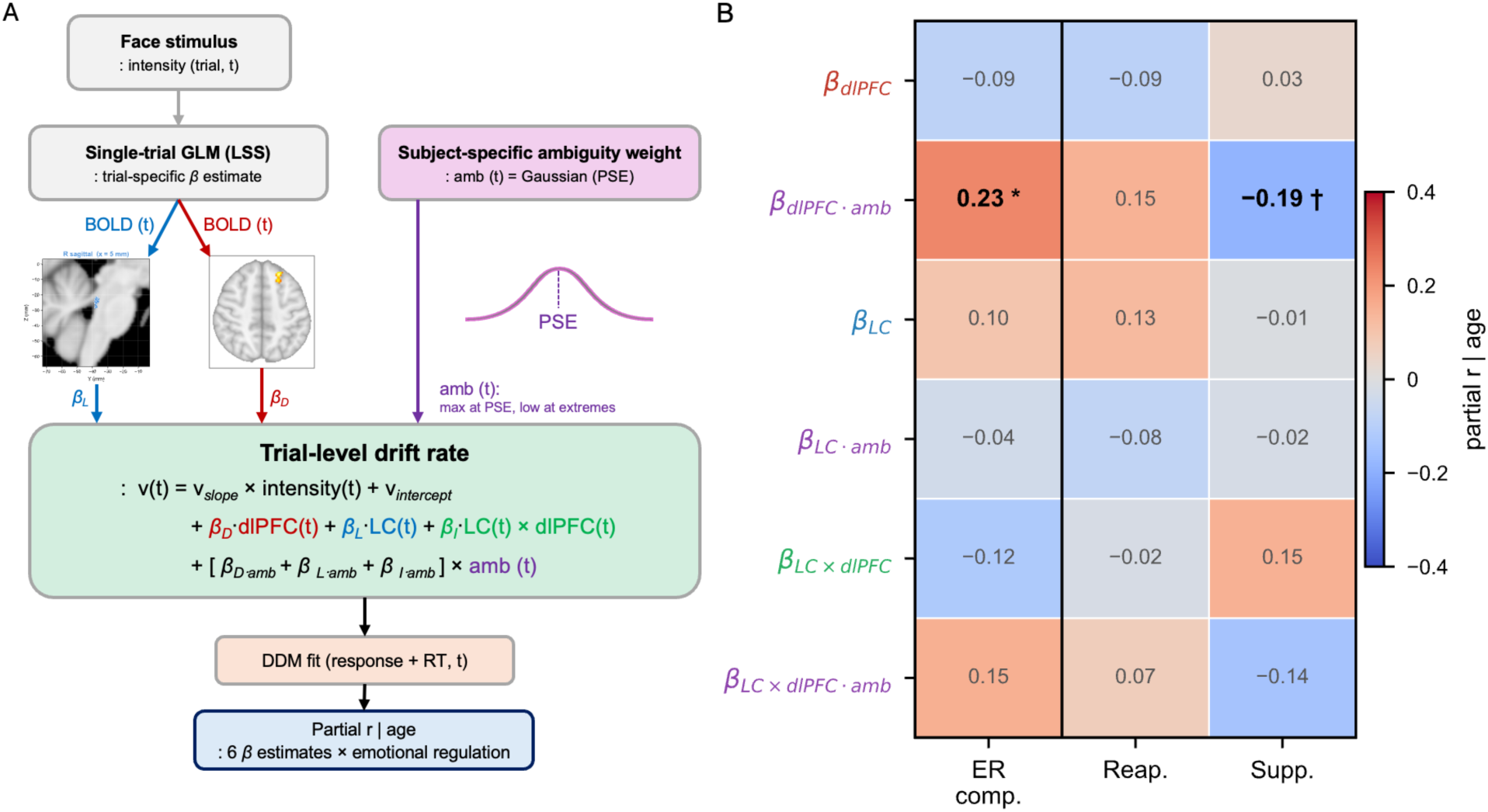
Trial-level DDM dissociates LC and dlPFC contributions to evaluative bias. (A) Model schematic. On each trial, single-trial BOLD was estimated using least-squares single-trial (LSS) modeling and extracted from two regions: the right LC peak voxel for each participant and the left dlPFC cluster defined by the group-level gPPI analysis. Trial-level drift rate was parameterized as a linear combination of stimulus intensity, LC activity (β_L_), dlPFC activity (β_D_), and their interaction (β_LC×dlPFC_), together with ambiguity-moderated variants of each term (amb), where trial-level ambiguity was defined as a subject-specific Gaussian weighting centered on each participant’s PSE, maximal at the PSE and minimal at the stimulus extremes. Six subject-level β parameters were estimated from each participant’s DDM fit across all trials. (B) Heatmap of age-controlled partial correlations between the six β parameters (rows) and three behavioral measures independent of the ROI definition (columns: cognitive reappraisal, expressive suppression, and ER composite). Row colors indicate the source term: red = dlPFC, blue = LC, green = LC × dlPFC interaction, and purple = ambiguity-moderated variant. The dlPFC ambiguity moderator (β_dlPFC·amb_) was associated with the ER composite (partial r | age = .23, p = .027), with a trend-level effect for suppression in the opposite direction (−.19, p = .08), whereas LC terms were uniformly null (all |r| < .13). Interaction terms were likewise null (all |r| < .15). This pattern indicates that dlPFC carried the individual-difference signal linked to emotion regulation, whereas a reliable LC contribution was not detected at either the group or individual-difference level. _†_*p* = .089 \**p* < .05, \*\**p* < .01.

LC single-trial activity (β_L_) showed a trend-level but nonsignificant main effect on drift (M=0.030, p=.096). LC terms were also not associated with individual differences in emotion regulation: age-controlled partial correlations between the LC terms (β_L_, β_L·amb_) and the behavioral measures were all smaller than |r|=.13 and none approached significance.

dlPFC activity (β_D_) also showed a trend-level main effect on drift (M=−.041, p=.053), but βD was not reliably associated with any emotion-regulation construct after controlling for age (ER composite: p=.418; reappraisal: p=.392; suppression: p=.761). By contrast, the ambiguity-moderated dlPFC term (β_D·amb_) showed a reliable age-controlled association with the ER composite (partial r | age=.234, 95% CI [.027,.422], p=.027), and a trend-level association in the opposite direction for expressive suppression (partial r | age=−.185, p=.08). β_D·amb_ was also numerically associated with positive bias (partial r | age=.279) and drift intercept (partial r | age=.221). These associations should be interpreted descriptively because the dlPFC cluster was originally identified through its between-person gPPI relationship with positive bias.

However, β_D·amb_ did not show a reliable group-level main effect on drift (p=.959), indicating that it indexed between-subject variability in ambiguity-dependent dlPFC coupling to drift rate rather than a uniform population-level shift. The LC × dlPFC interaction terms (β_I_, β_I·amb_) showed no reliable group-level effects or individual-difference associations with the behavioral measures. Because gPPI indexes task-related temporal covariation between LC and dlPFC, whereas β_D·amb_ captures the within-person mapping between single-trial dlPFC activity and drift rate near the participant’s perceptual boundary, the trial-level DDM result should be interpreted as a distinct but nonindependent signal. Given that the dlPFC ROI was defined from the primary positive-bias gPPI analysis, this pattern provides descriptive convergence within the bias-related LC–dlPFC circuit, not independent confirmatory evidence. LC activity, by contrast, did not show a reliable unique single-trial contribution when modeled simultaneously with dlPFC. The full pattern of trial-level associations is shown in **Fig 4B** and **Table S3.** Sensitivity analyses with a linear ambiguity-weighted trial-level DDM yielded the same pattern, see Supplementary Materials (**Fig S6**).

### LC temporal and structural properties relate to bias-defined cortical coupling

The trial-level analysis suggested that the behaviorally relevant ambiguity-dependent drift signal was expressed more clearly in dlPFC than in LC, raising the complementary question of whether LC was related to the bias-related circuit through more sustained signal properties. We therefore computed the Hurst exponent of each participant’s LC seed time series as an index of temporal persistence in the BOLD signal (N=90; M=0.517)

Because the dlPFC cluster was defined in the primary gPPI analysis by its association with positive bias, we treated the extracted LC–dlPFC coupling estimate as a bias-defined circuit phenotype rather than as an independent test of the coupling–bias association. Greater LC temporal persistence was associated with weaker, less negative LC–dlPFC coupling (partial r | age=.385, 95% CI [.194,.549], p<.001; **Fig 5A**) and, for interpretive reference, lower positive bias (partial r | age=−.246, 95% CI [−.434,−.045], p=.019; **Fig 5B**). By contrast, dlPFC temporal persistence (M=.647) was not reliably associated with either measure (all ps > .140), suggesting that the temporal-persistence association was more evident in the LC signal profile.

**Fig. 5.**
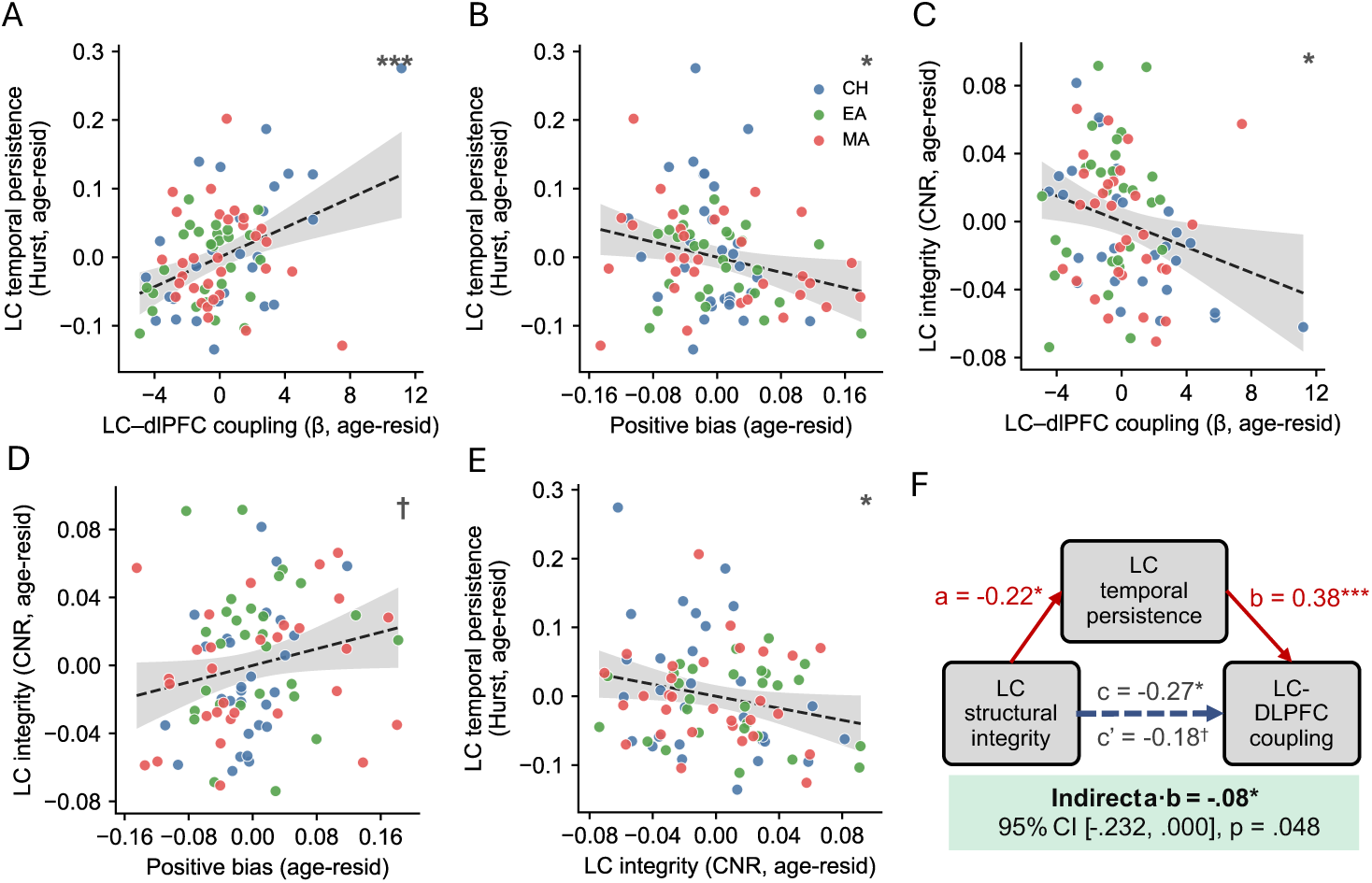
LC temporal persistence and neuromelanin-sensitive contrast are associated with bias-defined LC–dlPFC coupling. (A) ROI analysis of LC temporal persistence, indexed by Hurst values extracted from each participant’s LC seed time series, and LC–dlPFC coupling extracted from the bias-defined dlPFC cluster shown in Fig. 3A. Greater LC temporal persistence was associated with weaker, less negative LC–dlPFC coupling. (B) LC temporal persistence was also associated with lower positive bias. (C) ROI analysis of LC neuromelanin contrast-to-noise ratio (CNR), used as a proxy for LC structural integrity, and LC–dlPFC coupling extracted from the same cluster. Higher LC CNR was associated with stronger, more negative LC–dlPFC coupling. (D) LC CNR showed a trend-level positive association with positive bias. (E) Higher LC CNR was associated with lower LC temporal persistence. (F) Exploratory bootstrap mediation model testing whether LC temporal persistence statistically accounted for part of the association between LC CNR and LC–dlPFC coupling, controlling for age. The indirect path through LC temporal persistence was reliable but modest. Color coding: CH = children/adolescents, EA = emerging adults, MA = middle-aged adults. _†_*p* =.079, \**p < .05,**p < .01, \*\*\**p < .001.

LC neuromelanin-sensitive contrast, used here as a proxy for LC structural integrity, showed the opposite pattern. Higher LC CNR was associated with stronger, more negative LC–dlPFC coupling (partial r | age=−.262, 95% CI [−.448,−.054], p=.014; **Fig 5C**), lower LC temporal persistence (partial r | age=−.224, 95% CI [−.415,−.014], p=.037; **Fig 5E**), and a trend-level positive association with positive bias (partial r | age=.212, p=.051; **Fig 5D**). Thus, LC CNR and LC temporal persistence were related in opposite directions to the same bias-defined coupling phenotype.

Finally, an exploratory bootstrap mediation model tested whether LC temporal persistence statistically accounted for part of the association between LC CNR and LC–dlPFC coupling, controlling for age (N=87; 5,000 iterations; **Fig 5F**). The indirect effect was reliable but modest (a·b=−.084, 95% CI [−.232,−.0002], bootstrap p=.048; 31.5% mediated), and the direct effect of LC CNR on coupling was reduced from significant (c: β=−.27, p=.015) to marginal (c′: β=−.18, p=.08). Because Hurst and gPPI estimates were both derived from the LC seed time series, this model should be interpreted as a statistical decomposition of shared LC signal properties rather than evidence for a causal structural-to-functional pathway.

## DISCUSSION

The present study examined how emotionally ambiguous faces are resolved into categorical judgments across development and whether this process is linked to LC–dlPFC circuitry. Three findings emerged. First, participants became less likely to resolve ambiguous angry-to-happy faces negatively across development. This age-related increase in positive interpretation was captured by converging SDT and DDM indices. The effect was not explained by perceptual sensitivity, starting point, response caution, or nondecision time, indicating that positive ambiguity resolution reflected a stimulus-independent evaluative drift bias. Second, this evaluative-bias dimension was linked to right LC–left dlPFC coupling during each participant’s subject-specific ambiguity condition. Participants with greater positive bias showed more negative task-modulated LC–dlPFC gPPI estimates. Third, trial-level modeling suggested that ambiguity-dependent drift was expressed more clearly in dlPFC than in LC, whereas LC temporal persistence and structural integration were related to the bias-defined LC–dlPFC coupling phenotype. Together, these findings suggest that developmental differences in emotional ambiguity resolution reflect evaluative evidence accumulation supported by LC–dlPFC circuitry.

### Positive Bias as Reduced Negative Evaluative Drift

Developmental increases in positive interpretation reflected reduced negative evaluative drift, not greater perceptual sensitivity or a general positive response tendency. Across groups, mean PSE values were at or above the morph midpoint, and drift intercepts were negative in children and emerging adults but approached zero in middle-aged adults. This pattern suggests a weakening of the negative default that often characterizes ambiguous facial-expression perception, rather than a uniform tendency to perceive ambiguous faces as happy (Petro et al., 2018). In this sense, the developmental effect is better described as reduced negative evaluative drift than as increased positivity per se.

The DDM results localized this shift to evaluative drift. Positive bias was not explained by sharper perceptual discrimination, because neither psychometric sensitivity nor drift slope accounted for the effect. It was also not explained by pre-evidence response bias or response caution, because starting point and boundary separation did not track positive bias. Instead, positive interpretation was captured specifically by drift intercept, a stimulus-independent component of evidence accumulation. Thus, the model-based results distinguish evaluative drift from perceptual gain, prior response bias, and decision threshold.

The parameter-specific connectivity analyses reinforced this interpretation. When the whole-brain analysis was repeated with alternative SDT and DDM covariates, drift intercept was the only decision parameter that recovered the left dlPFC cluster identified by positive bias. Nondecision time produced medial frontal, cingulate, and parietal LC connectivity patterns, but these effects are more consistent with timing, response preparation, or task-readiness processes than with ambiguity resolution. Thus, the LC–dlPFC effect was not a generic marker of task difficulty, decision speed, or model-wide performance variation. It was most closely aligned with the evaluative drift component of ambiguity resolution.

This finding builds directly on the recent top-down framework of emotional perception articulated by Mohanty et al. (2025), which provides an important synthesis of how prior knowledge, attention, context, and evaluative goals shape affective perception. The present results add process-level specificity to that framework by showing that, in this developmental sample, positive ambiguity resolution was expressed as reduced negative evaluative drift rather than perceptual gain or starting-point bias.

### Subjective Ambiguity and LC–dlPFC Coupling

Ambiguity in emotional perception should be treated as an observer-relative decision state, not only as a fixed stimulus property. In the present sample, the morph condition closest to each participant’s PSE often differed from the nominal neutral midpoint. Consistent with this view, the LC–dlPFC connectivity effect emerged when analyses were anchored to each participant’s PSE-nearest ambiguity level, but not when ambiguity was defined by a fixed neutral condition or by a stimulus-defined high-ambiguity average. This pattern suggests that LC–dlPFC coupling was linked to the participant’s actual point of categorical uncertainty, where evaluative bias should be most consequential.

### Relation to Prior LC–Prefrontal Findings

The LC–dlPFC cluster observed here is anatomically close to the prefrontal region implicated by Dave et al., who found increased LC activity and stronger LC–prefrontal coupling during ambiguity processing in older adults (Dave et al., 2025). The present findings are therefore best viewed as a developmental extension rather than a direct replication. Whereas that work established LC–prefrontal coupling as a candidate ambiguity-resolution circuit in older adulthood, the present study extends this circuit to late childhood through middle adulthood and links it specifically to evaluative drift bias.

The difference in coupling-bias direction should be interpreted cautiously. gPPI estimates reflect task-modulated coupling relative to an implicit baseline, so a negative estimate does not necessarily indicate weaker absolute connectivity. The two studies also defined ambiguity differently: the present study used participant-specific PSE-nearest ambiguity, whereas Dave et al. used a fixed stimulus-defined contrast. Finally, the studies sampled different age ranges. Thus, the conservative interpretation is that both studies implicate LC–prefrontal circuitry in affective ambiguity resolution, while suggesting that its functional expression may differ by developmental context and ambiguity definition.

### Emotion Regulation as a dlPFC Anchor

Emotion-regulation measures were included to anchor the proposed dlPFC role in evaluative ambiguity resolution. Reappraisal is theoretically relevant because it depends on prefrontal systems that reinterpret emotional meaning, whereas suppression acts later on expressive behavior. Consistent with this distinction, the ER composite was associated with positive bias at the SDT level and showed a marginal association with drift intercept. Cognitive reappraisal was also linked to the bias-defined LC–dlPFC coupling phenotype, whereas expressive suppression was not. This pattern fits the idea that the dlPFC component of the LC–dlPFC circuit is related to evaluative reinterpretation rather than to response inhibition or general regulation tendency.

The trial-level DDM further supported this framing. The ambiguity-moderated dlPFC term, rather than the LC terms, carried the clearest individual-difference signal linked to emotion regulation. Thus, the ER findings are best interpreted as converging evidence that dlPFC contributes to evaluative reinterpretation near the participant’s perceptual boundary, not as evidence that habitual reappraisal directly causes positive ambiguity resolution.

### Trial-Locked dlPFC Evaluation and Sustained LC Gain

The neural results suggest a timescale dissociation: dlPFC carried the clearer trial-locked evaluative signal, whereas LC was related to sustained signal properties of the bias-related circuit. gPPI identified LC–dlPFC coupling as a between-person circuit phenotype, but it cannot determine whether LC, dlPFC, or their joint activity carries trial-level decision information. When LC and dlPFC activity were entered simultaneously as predictors of drift rate, the clearer individual-difference signal was found in the ambiguity-moderated dlPFC term. LC terms, by contrast, showed no reliable unique trial-level contribution when modeled together with dlPFC. This trial-level result is therefore best viewed as a within-circuit follow-up rather than an independent localization of a new effect.

This pattern is consistent with adaptive-gain accounts of LC function, in which LC–norepinephrine activity regulates the balance between temporally focused task engagement and broader tonic arousal state (Aston-Jones & Cohen, 2005; Mather et al., 2016; Sara, 2009). In this framework, dlPFC may express the trial-locked evaluative signal that shapes evidence accumulation on individual ambiguous trials, whereas LC may contribute through a sustained, tonic-like gain state that shapes how strongly cortical evaluative signals are coupled to task context. LC temporal persistence, indexed by the Hurst exponent, provided an operational measure of this sustained BOLD signal profile, consistent with prior work treating temporal variability and persistence in BOLD activity as meaningful individual-difference features (Garrett et al., 2013): it was associated with the bias-defined LC–dlPFC coupling phenotype, whereas dlPFC temporal persistence was not. Higher LC neuromelanin-sensitive CNR, used here as a proxy for LC structural integrity (Clewett et al., 2016; Neal et al., 2025), was also associated with lower LC temporal persistence and with stronger, more negative LC–dlPFC coupling

This interpretation remains indirect. Hurst indexes temporal persistence in the BOLD signal, not tonic LC firing directly, and neuromelanin-sensitive CNR is a proxy for LC structural integrity rather than a direct measure of noradrenergic cell density or neurotransmitter release. Because Hurst and gPPI estimates were derived from the same LC seed time series, the mediation analysis should be treated as an exploratory statistical decomposition of shared LC signal properties rather than evidence for a causal structural-to-functional pathway.

### Limitation and Conclusion

Several limitations constrain these conclusions. The cross-sectional design and lack of older adults limit claims about within-person developmental change or direct convergence with Dave et al. (Dave et al., 2025). gPPI cannot establish directional influence between LC and dlPFC. Trial-level LC estimates came from small brainstem signals at 2.5-mm functional resolution, so null LC effects should be interpreted cautiously. Finally, the task used an angry-to-happy continuum, and whether the same evaluative drift mechanism applies to other forms of ambiguity remains unknown.

In sum, positive interpretation of emotional ambiguity was best characterized as reduced negative evaluative drift rather than as increased perceptual sensitivity, starting-point bias, response caution, or general task performance. This evaluative-bias dimension was linked to reappraisal, ambiguity-dependent dlPFC activity, and task-modulated LC–dlPFC coupling. The findings extend LC–prefrontal ambiguity-resolution circuitry to earlier development and identify evaluative drift bias as a decision-process signature of emotional ambiguity resolution.

## Author Contributions

THL and YY Chen developed the study concept. THL analyzed and interpreted data. THL and YYC drafted the manuscript, and THL, YYC, and JN provided critical revisions. All authors approved the final version of the manuscript for submission.

## Acknowledgements

This work was done based on research supported by R01AG075000/AG/NIA NIH and by 2022 Virginia Tech Lay Nam Chang Dean’s Discovery Fund. This work was also supported by Virginia Tech Institute for Society, Culture and Environment research award. In preparing this manuscript, the authors used *Claude* for language editing, sentence-level revision and for assistance with code generation used to render figure layouts. All data analyses, scientific content, interpretations, and conclusions were developed by the authors, who reviewed and approved the final manuscript and take full responsibility for its content. All scientific content, interpretations, and conclusions were developed by the authors, who reviewed and approved the final manuscript and take full responsibility for its content.

## SUPPLEMENTARY MATERIALS

### Subject-Specific Maximum-Ambiguity Definition

To account for individual differences in the perception of emotional ambiguity, we defined subject-specific ambiguity levels based on each participant’s PSE estimated above using logistic fit. The continuous PSE value was linearly transformed to a scale ranging from −3 to +3 using the formula: scale = (PSE − 0.5) × 6. This transformed value was then mapped onto the nearest of the seven discrete morph conditions (AN60 = −3, AN35 = −2, AN15 = −1, NE = 0, HA15 = 1, HA35 = 2, HA60 = 3), yielding a participant-specific PSE-nearest morph condition. The condition closest to each participant’s transformed PSE value was treated as that individual’s point of maximum subjective ambiguity, that is, the morph level at which angry and happy judgments were equally likely. For example, a participant with a PSE of 0.60 would have a transformed score of 0.60 and would therefore be assigned to the nearest discrete condition, HA15, which represented that participant’s maximum-ambiguity condition. In the present sample (N = 90), PSE-nearest morph conditions were distributed as follows: HA15 (n = 52, 57.8%), HA35 (n = 15, 16.7%), NE (n = 10, 11.1%), AN15 (n = 9, 10.0%), and AN35 (n = 4, 4.4%); no participant’s PSE-nearest condition fell at AN60 or HA60, consistent with the overall tendency to interpret ambiguous faces as somewhat negative (group-mean PSE > 0.5 on the [0, 1] morph scale). In all subsequent group-level analyses, including gPPI and task-evoked GLM analyses, we extracted each participant’s first-level contrast of parameter estimates (cope) corresponding to the participant-specific PSE-nearest morph condition rather than using a fixed, stimulus-defined ambiguity midpoint. This approach ensured that connectivity estimates reflected each participant’s subjective experience of the highest ambiguity rather than stimulus-defined category boundaries that might not align with individual perceptual thresholds (**Fig S1**).

To verify that the left dlPFC cluster reflected subject-specific rather than stimulus-defined ambiguity processing, we re-ran the same group-level analysis using two alternative contrast definitions. First, we used a fixed neutral-midpoint cope (NE) for every participant (Dave et al., 2025). Second, we averaged the three highest-ambiguity conditions (NE, AN15, and HA15) into a single high-ambiguity cope for each participant (Wang et al., 2017).

Under both fixed-contrast schemes, no cluster in the left dlPFC survived correction at any LC seed definition. Our main finding (LC-dlPFC) therefore depended specifically on sampling each participant’s gPPI at that participant’s own maximum-ambiguity condition, supporting the interpretation that this cluster reflects subject-specific interpretive processing rather than stimulus-locked features of the morph midpoint. The right angular gyrus cluster, in contrast, generalized across subject-specific and stimulus-defined sampling schemes, consistent with the possibility that this parietal effect reflects a less subject-specific component of ambiguity processing.

**Fig S1.**
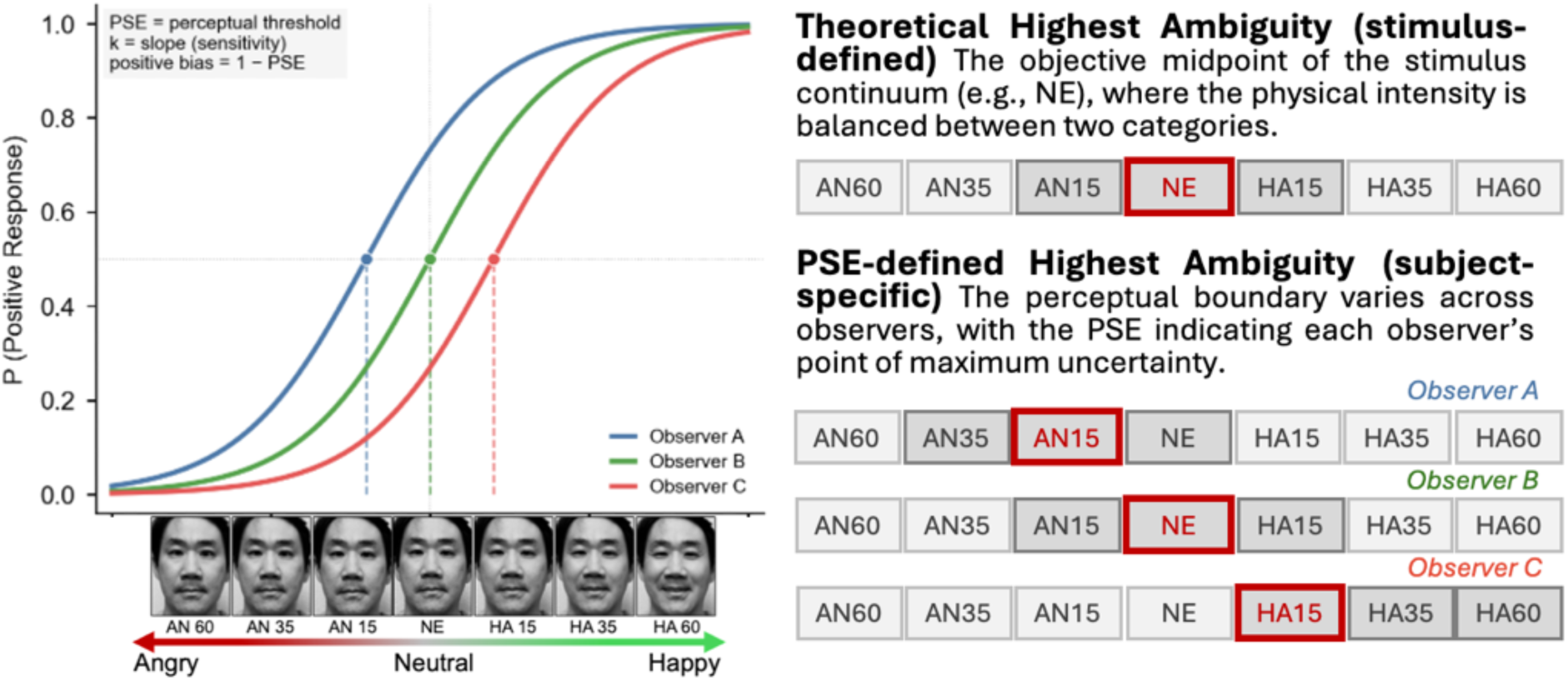
Schematic illustration of PSE-individualized ambiguity assignment. Logistic psychometric functions were fitted to each participant’s behavioral data, and the point of subjective equality (PSE) was defined as the morph level at which the participant was equally likely to categorize a face as angry or happy. Colored curves illustrate three example participants with low (red), intermediate (green), and high (blue) PSE values, whereas gray curves show the remaining participants. For visualization, the example curves are shown with a common slope. (Right) Comparison of stimulus-defined and PSE-defined ambiguity classification. In the stimulus-defined approach, the highest-ambiguity condition is fixed across observers. In the PSE-defined approach, the condition closest to each participant’s PSE is designated as the highest ambiguity condition (red boxes), with adjacent conditions classified as lower ambiguity. Note. Photographs shown in the task schematic are of the authors and are reproduced with their permission.

### Robustness of LC seed definition

Because LC is an exceptionally small midbrain nucleus (sensitive to motion and other artifacts) and the seed used in the primary analysis was centered on each participant’s neuromelanin-defined LC peak voxel, we conducted an additional sensitivity analysis in which the seed sphere was systematically expanded around the same peak. The left dlPFC cluster replicated across all right-sided LC seed spheres (2, 3, and 4 mm radius), with cluster size, at corrected p < .05, increasing monotonically with seed size: 245 voxels for the 2 mm sphere, 289 voxels for the 3 mm sphere, and 312 voxels for the 4 mm sphere, as well as 172 voxels for the atlas-based LC seed. In every case, greater positive bias was associated with weaker, more negative LC–dlPFC connectivity, and no cluster survived correction under the opposite contrast. This pattern confirms that the observed LC–dlPFC relationship was unidirectional and robust both to seed size and to seed-localization strategy. The right angular gyrus finding also replicated across all left LC seed sphere sizes (2 mm sphere: 228 voxels; 3 mm sphere: 215 voxels; 4 mm sphere: 198 voxels), as well as with the probabilistic LC atlas seed (344 voxels), confirming that this secondary finding was likewise robust to seed definition.

**Fig S2.**
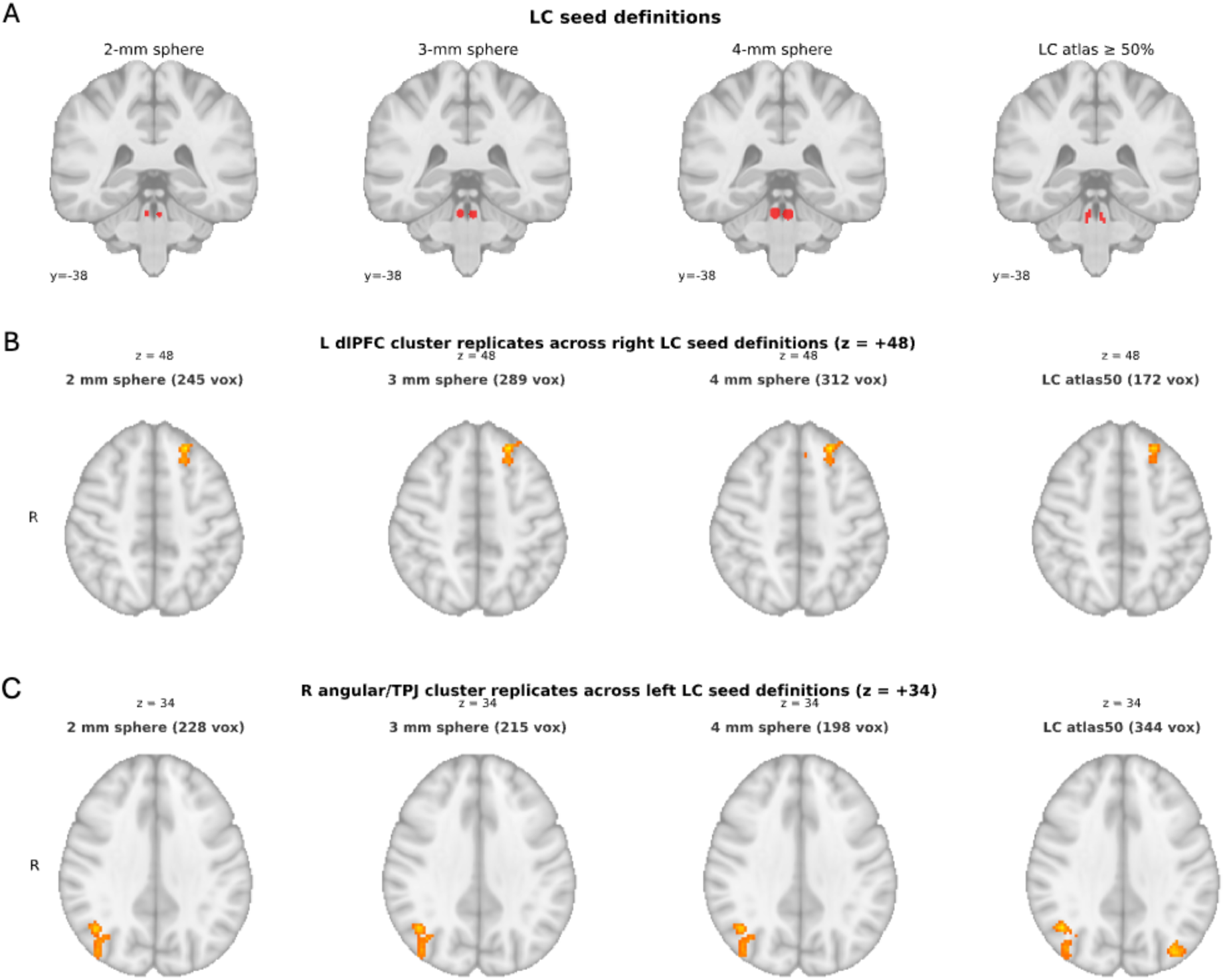
Robustness of LC seed definition. Four LC seed variants used to verify that the gPPI results were not driven by a choice of seed extent. From left to right, the figure shows 2 mm (primary seed in the main draft), 3 mm, and 4 mm spheres centered on each participant’s LC peak voxel, with the group-mean peak shown here, R LC at 5, −38, −24 and L LC at −4, −38, −25, as well as the probabilistic LC atlas thresholded at 50%. Masks are overlaid on the MNI152 1 mm template at the coronal slice y = −38 in radiological orientation. Whole-brain gPPI analyses relating connectivity to positive bias were repeated using each of these seed definitions. (B) Whole-brain gPPI × positive-bias contrast (z > 2.57, cluster-corrected p < .05) for the right-LC seed across the four definitions. Each analysis recovered the same left dlPFC cluster reported in the main text, indicating that the R LC → L dlPFC coupling effect is not dependent on the choice of seed radius or on the use of subject-specific peak-voxel seeds versus the atlas-based seed. (C) Matching whole-brain contrast for the left-LC seed, which in every seed definition recovered the same right angular gyrus / TPJ cluster. Together these panels show that the primary connectivity findings are stable across seed size and across peak-voxel versus atlas-based seed construction.

### Whole-brain Estimation with Other Parameters

To test whether the LC–dlPFC circuit was specific to the evaluative-bias dimension or instead reflected a broader association with perceptual decision-making, we repeated the group-level gPPI analysis while substituting each SDT and DDM parameter for positive bias as the behavioral covariate, with age included as a covariate.

Of the seven candidate parameters (SDT: sensitivity k; DDM: drift intercept, drift slope, boundary separation, starting point, and non-decision time; plus ER composite), only drift intercept yielded a reliable cluster. When drift intercept was entered as the covariate, the negative-coupling contrast again revealed a left dlPFC cluster (peak −20, 34, 50; 118 voxels; p = .027; **Fig S3**) whose location was essentially identical to that of the positive-bias cluster (peak −22, 34, 48), confirming that the same LC–dlPFC circuit captured the bias-related construct whether it was indexed by perceptual threshold (1 − PSE) or by the stimulus-independent component of evidence accumulation (drift intercept). Non-decision time also produced significant right LC connectivity clusters in medial frontal/cingulate and parietal regions. Because non-decision time captures non-evaluative aspects of performance, including encoding, response preparation, and motor execution, we interpreted these clusters as reflecting general timing or task-readiness processes rather than positive interpretive bias. We therefore report this result as part of the specificity analyses but do not pursue it as a primary ambiguity-resolution circuit. By contrast, perceptual sensitivity (slope k), drift slope, boundary separation, starting point, and ER composite produced no suprathreshold clusters under any LC seed definition. The null results for sensitivity and drift slope are consistent with the behavioral dissociation reported above: LC–cortical coupling tracked the content of evaluative bias, rather than perceptual gain. The null result for starting point is likewise consistent with the observation that x₀ carried little behavioral bias variance (r = .081, n.s.), reinforcing the conclusion that positive interpretive bias is implemented at the level of drift rather than as a pre-evidence expectation prior. We note that no family-wise correction was applied across the seven covariates tested; the specificity claim rests on the overall pattern of results, in which only drift intercept recovered the dlPFC cluster, rather than on formal multiplicity control across covariates.

**Fig. S3.**
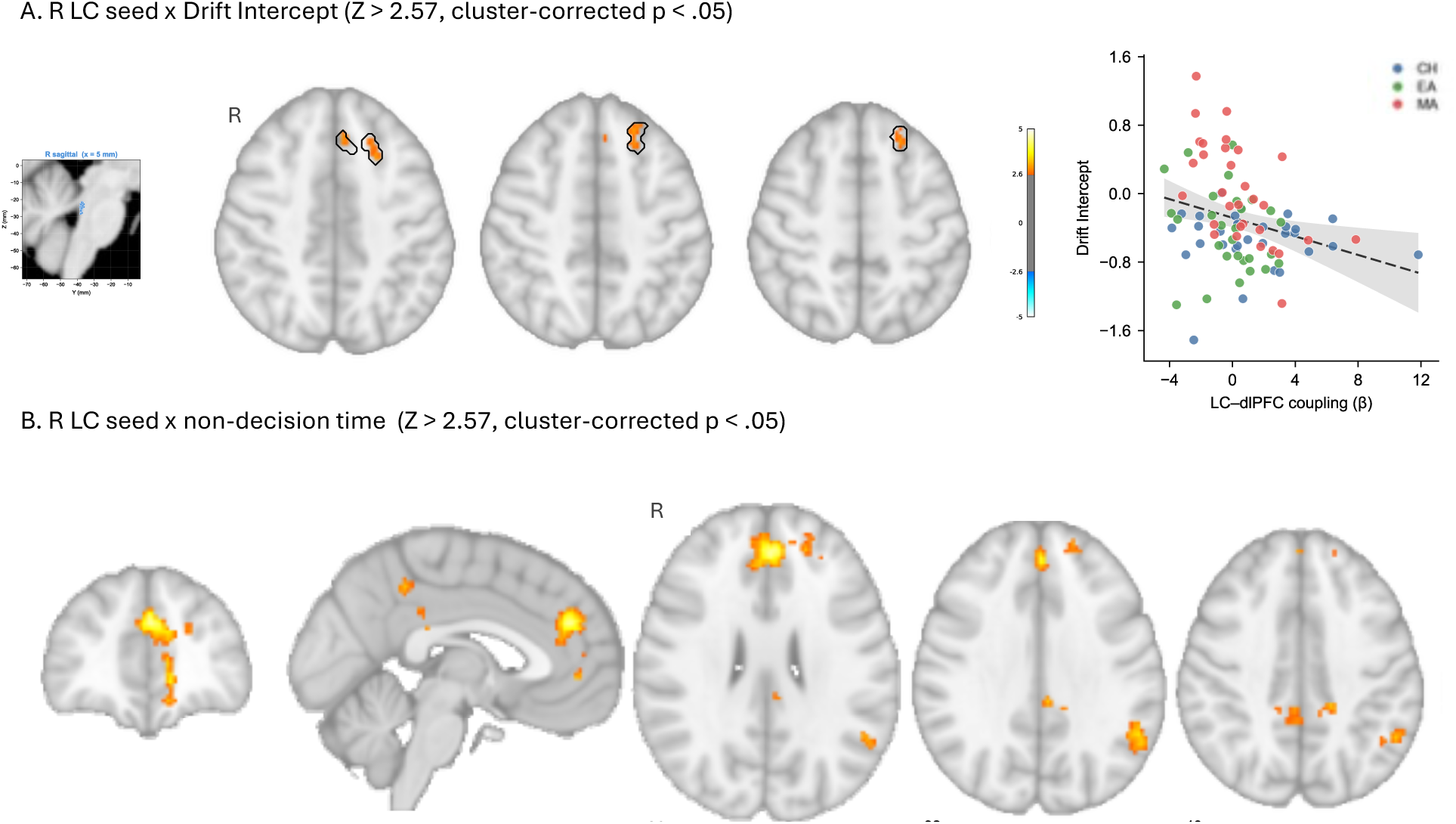
LC–dlPFC connectivity associated with DDM parameters. (A) Representative right LC seed location, whole-brain gPPI result, and extracted ROI scatterplot. In the right LC seed analysis, with drift intercept entered as the behavioral covariate and age included as a nuisance covariate, higher drift intercept was associated with lower right LC connectivity estimates in a left dlPFC cluster (peak MNI: −20, 34, 50; 118 voxels; FLAME 1+2, Z > 2.57, cluster-corrected p < .05). This cluster closely overlapped the positive-bias cluster shown in Fig. 3A (peak MNI: −22, 34, 48; 245 voxels). The scatterplot is shown for visualization and was not used as an independent inferential test. (B) Right LC seed analysis with nondecision time entered as the behavioral covariate, controlling for age. Nondecision time was associated with LC connectivity in medial frontal/cingulate and parietal regions, including medial superior frontal cortex, pregenual ACC, precuneus, mid/posterior cingulate, inferior parietal cortex, supramarginal gyrus, and angular gyrus. Because nondecision time reflects non-evaluative components of task performance, including stimulus encoding, response preparation, and motor execution, these clusters were interpreted as a general timing or task-readiness-related LC connectivity pattern rather than as part of the positive-bias ambiguity-resolution pathway.

**Table S1.**
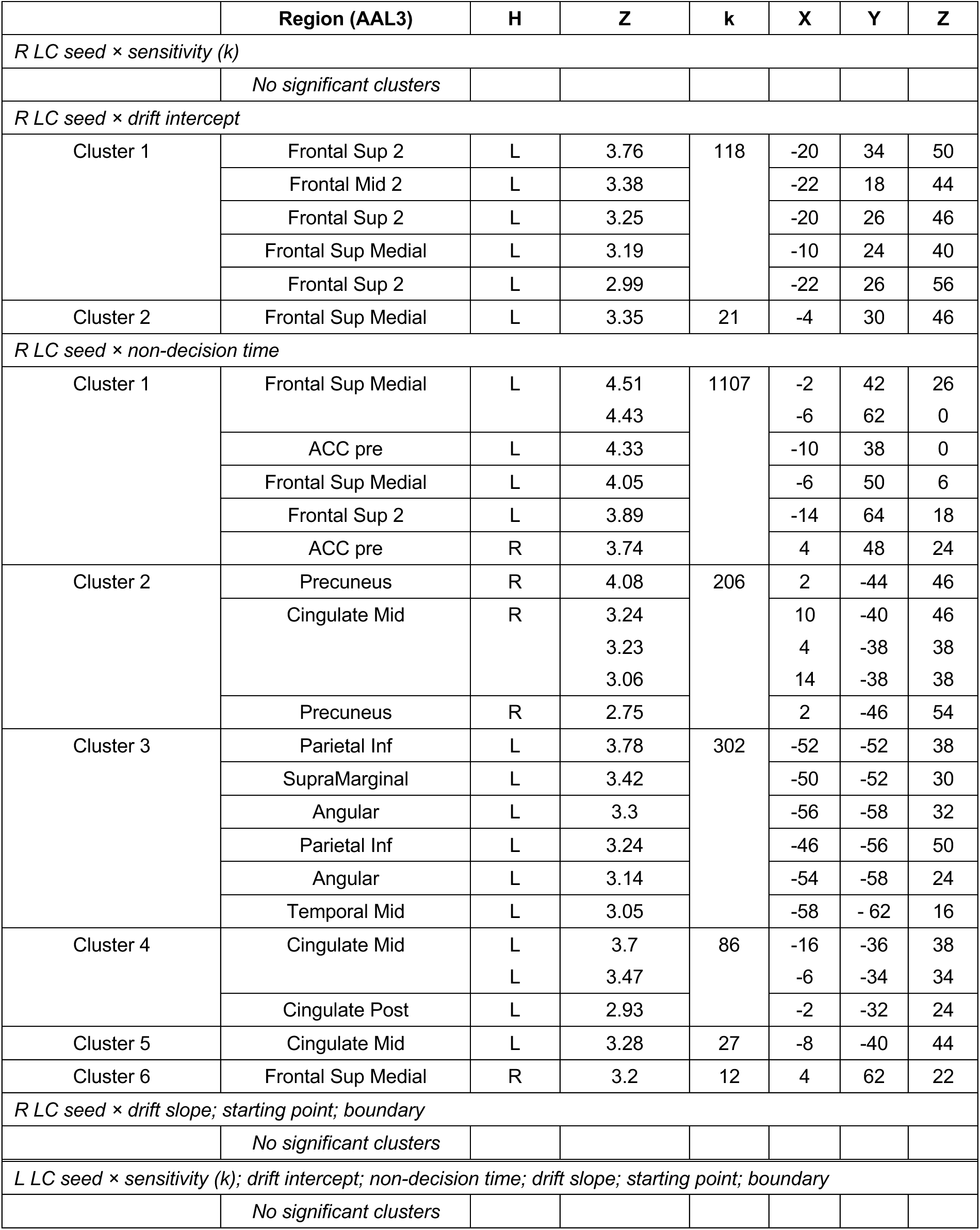
Whole-brain gPPI parameter follow-up: LC connectivity as a function of behavioral and decision-model parameters. To test whether the LC–dlPFC circuit identified in the primary analysis was specific to the evaluative-bias dimension, the group-level gPPI analysis was repeated with each candidate covariate substituted for positive bias, with age included as a covariate (FLAME 1+2, Z > 2.57, cluster-corrected p < .05). Regions were labeled using the AAL3v1 atlas. Of the candidate covariates tested, only drift intercept recovered the left dlPFC cluster, supporting the interpretation that LC–cortical coupling was most closely tied to the evaluative-bias construct. H = hemisphere; Z = peak Z statistic; k = cluster size in 2-mm isotropic voxels; x, y, z = MNI coordinates in millimeters.

|  | Region (AAL3) | H | Z | k | X | Y | Z |
| --- | --- | --- | --- | --- | --- | --- | --- |
| *R LC seed × sensitivity (k)* |  |  |  |  |  |  |  |
|  | No significant clusters |  |  |  |  |  |  |
| *R LC seed × drift intercept* |  |  |  |  |  |  |  |
| Cluster 1 | Frontal Sup 2 | L | 3.76 | 118 | -20 | 34 | 50 |
|  | Frontal Mid 2 | L | 3.38 |  | -22 | 18 | 44 |
|  | Frontal Sup 2 | L | 3.25 |  | -20 | 26 | 46 |
|  | Frontal Sup Medial | L | 3.19 |  | -10 | 24 | 40 |
|  | Frontal Sup 2 | L | 2.99 |  | -22 | 26 | 56 |
| Cluster 2 | Frontal Sup Medial | L | 3.35 | 21 | -4 | 30 | 46 |
| *R LC seed × non-decision time* |  |  |  |  |  |  |  |
| Cluster 1 | Frontal Sup Medial | L | 4.51 | 1107 | -2 | 42 | 26 |
|  |  |  | 4.43 |  | -6 | 62 | 0 |
|  | ACC pre | L | 4.33 |  | -10 | 38 | 0 |
|  | Frontal Sup Medial | L | 4.05 |  | -6 | 50 | 6 |
|  | Frontal Sup 2 | L | 3.89 |  | -14 | 64 | 18 |
|  | ACC pre | R | 3.74 |  | 4 | 48 | 24 |
| Cluster 2 | Precuneus | R | 4.08 | 206 | 2 | -44 | 46 |
|  | Cingulate Mid | R | 3.24 |  | 10 | -40 | 46 |
|  |  |  | 3.23 |  | 4 | -38 | 38 |
|  |  |  | 3.06 |  | 14 | -38 | 38 |
|  | Precuneus | R | 2.75 |  | 2 | -46 | 54 |
| Cluster 3 | Parietal Inf | L | 3.78 | 302 | -52 | -52 | 38 |
|  | SupraMarginal | L | 3.42 |  | -50 | -52 | 30 |
|  | Angular | L | 3.3 |  | -56 | -58 | 32 |
|  | Parietal Inf | L | 3.24 |  | -46 | -56 | 50 |
|  | Angular | L | 3.14 |  | -54 | -58 | 24 |
|  | Temporal Mid | L | 3.05 |  | -58 | -62 | 16 |
| Cluster 4 | Cingulate Mid | L | 3.7 | 86 | -16 | -36 | 38 |
|  |  | L | 3.47 |  | -6 | -34 | 34 |
|  | Cingulate Post | L | 2.93 |  | -2 | -32 | 24 |
| Cluster 5 | Cingulate Mid | L | 3.28 | 27 | -8 | -40 | 44 |
| Cluster 6 | Frontal Sup Medial | R | 3.2 | 12 | 4 | 62 | 22 |
| *R LC seed × drift slope; starting point; boundary* |  |  |  |  |  |  |  |
|  | No significant clusters |  |  |  |  |  |  |
| *L LC seed × sensitivity (k); drift intercept; non-decision time; drift slope; starting point; boundary* |  |  |  |  |  |  |  |
|  | No significant clusters |  |  |  |  |  |  |

**Fig S4.**
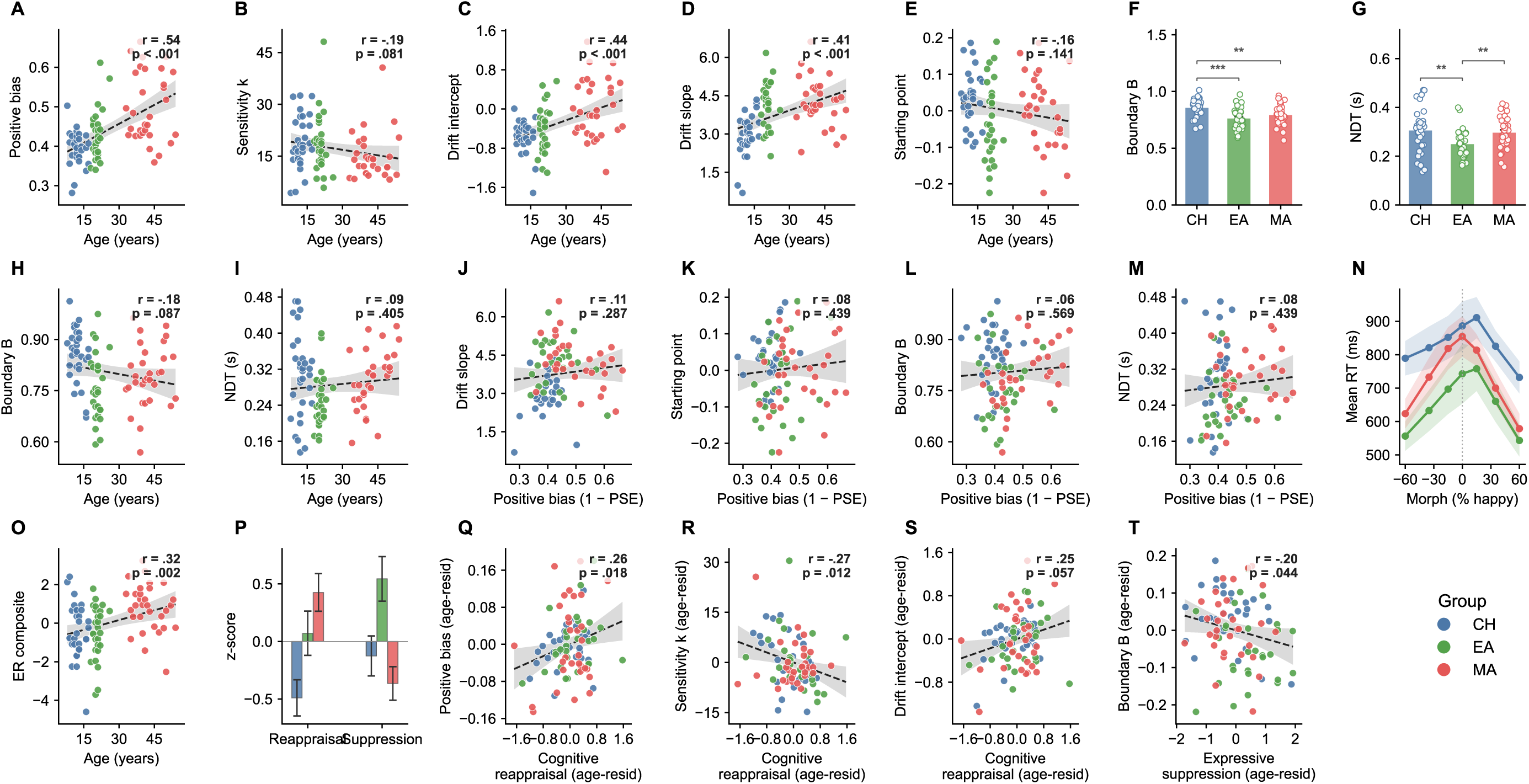
SDT and DDM parameter development and emotion-regulation subscale associations. **Row 1 (A–G).** Continuous age associations and group comparisons for SDT and DDM parameters. (A) Positive bias, indexed as 1 − PSE, increased with age. (B) Sensitivity, indexed by slope *k*, showed a weak negative association with age. (C) Drift intercept increased with age, paralleling the age effect on positive bias. (D) Drift slope increased with age. (E) Starting point did not show a reliable age association. (F) Boundary separation differed by developmental group, with children showing greater boundary separation than both adult groups. (G) Non-decision time differed by developmental group, with emerging adults showing the shortest non-decision times, consistent with a U-shaped developmental pattern. **Row 2 (H–N).** Remaining DDM parameter age associations, response-time profile, and specificity of positive bias across DDM components. (H) Boundary separation as a function of age, showing a weak linear decline across the full sample. (I) Nondecision time as a function of age, showing no monotonic linear age trend. (J–M) Specificity of positive bias across DDM parameters: drift slope (J), starting point (K), boundary separation (L), and nondecision time (M), each plotted against positive bias. Of the five DDM parameters, only drift intercept tracked positive bias at the individual level; the other four parameters shown here were not reliably associated with positive bias. Together, these panels show that boundary separation and nondecision time followed theoretically expected developmental patterns, but neither parameter explained individual differences in positive bias. (N) Mean reaction time (ms) by morph intensity and developmental group, with shaded 95% confidence bands. **Row 3 (O–T).** Emotion-regulation subscale associations with SDT and DDM components. (O) ER composite, computed as *z*(reappraisal) − *z*(suppression), as a function of age. (P) Cognitive reappraisal and expressive suppression by developmental group, expressed as z-scores. (Q) Cognitive reappraisal was positively associated with positive bias after controlling for age. (R) Cognitive reappraisal was negatively associated with perceptual sensitivity *k* after controlling for age. (S) Cognitive reappraisal showed a marginal positive association with drift intercept after controlling for age. (T) Expressive suppression was negatively associated with boundary separation after controlling for age. Color coding: CH, children/adolescents, blue; EA, emerging adults, green; MA, middle-aged adults, red. Scatter panels show raw values in Rows 1–2, panels A–E and H–N, and age-residualized values in Row 3, panels Q–T. Each scatter plot reports Pearson’s *r* and its *p* value. Dashed lines indicate linear fits with 95% confidence bands. Bar panels show group means ± SEM with individual data points overlaid. \**p* < .05, \*\**p* < .01, \*\*\**p* < .001.

### Whole-brain gPPI lightbox views for LC connectivity and post hoc ROI analysis of age-group patterns

The primary whole-brain finding was a negative association between right LC connectivity with left dlPFC and positive bias, as described in the main text. **Fig S5** provides lightbox views of the suprathreshold clusters from the whole-brain gPPI analyses to show the spatial extent of the LC connectivity patterns associated with positive bias. In addition to the right LC–left dlPFC finding, the left LC seed identified a right angular gyrus/temporoparietal junction cluster whose connectivity estimates were also negatively associated with positive bias, see **Table S2.**

We next conducted post hoc ROI analyses to characterize whether these connectivity–bias associations differed across age groups. For the primary LC–dlPFC pathway **(Fig S5C)**, we extracted each participant’s mean gPPI connectivity estimate from the left dlPFC cluster identified in the whole-brain analysis. We then examined both the mean connectivity estimate and its association with positive bias within each age group. The mean LC–dlPFC connectivity estimate did not vary linearly with age, p = .842, and did not show a significant group effect, p = .107. Instead, the age-group pattern was expressed in the strength of the connectivity–bias association. The association was small and nonsignificant in children, p = .13, marginal in emerging adults, r = −.41, 95% CI [−.75, .05], p = .076, and strongest in middle-aged adults, r = −.61, 95% CI [−.77, −.39], p < .001. Thus, the whole-brain LC–dlPFC effect was not explained by age-related differences in mean connectivity. Rather, the inverse association between LC–dlPFC connectivity and positive bias was most evident in middle-aged adults and weaker in the younger groups.

**Fig S5.**
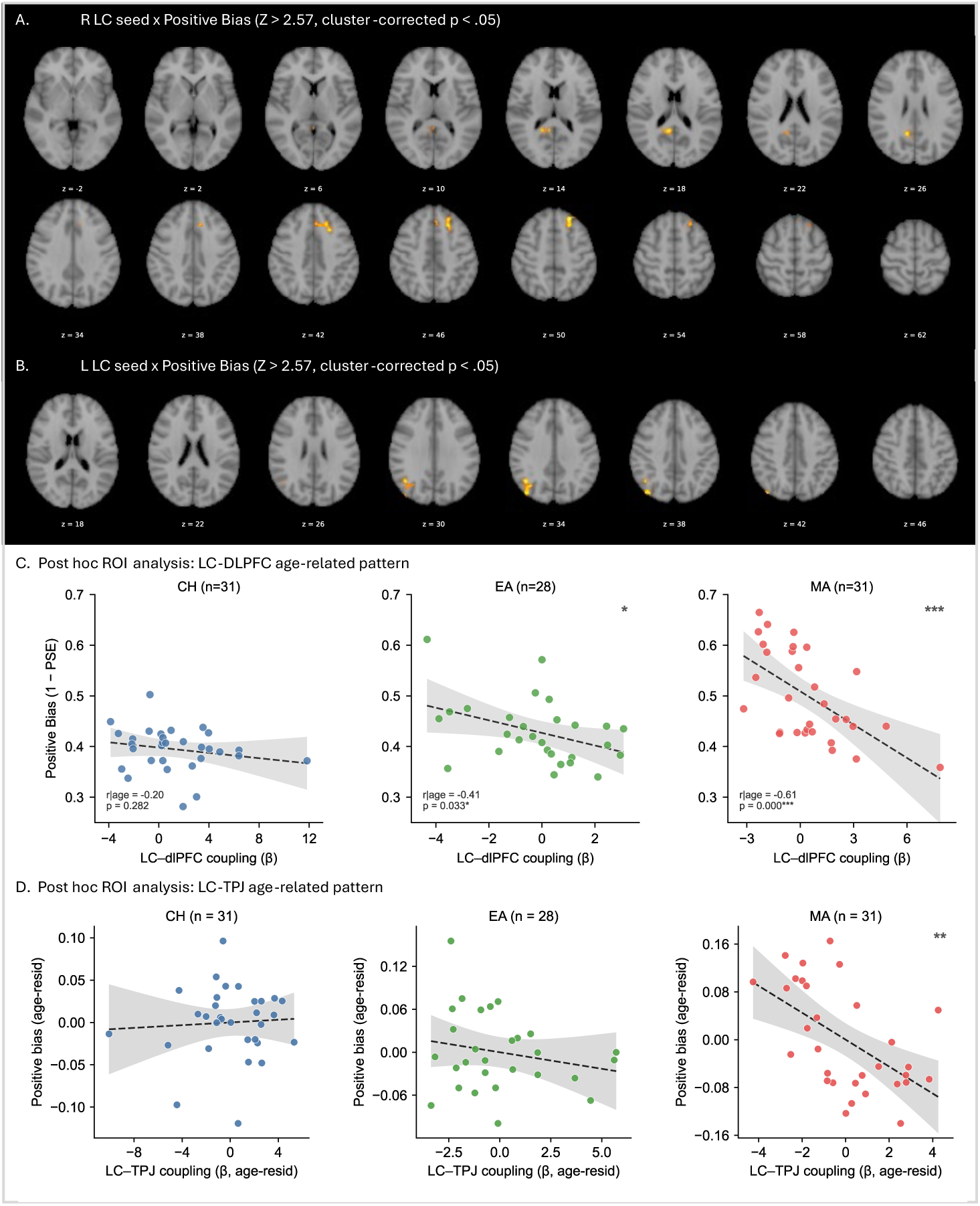
Whole-brain gPPI lightbox views and post hoc ROI age-group patterns for bias-related LC connectivity. (A) Right LC seed × positive bias, negative association. Axial slices containing suprathreshold voxels are shown, Z > 2.57, cluster-corrected p < .05, N = 90. The primary left dlPFC cluster, peak MNI: −22, 34, 48; 245 voxels, is visible across z = 34 to 54. (B) Left LC seed × positive bias, negative association. The secondary right angular gyrus/TPJ cluster, peak MNI: 44, −60, 32; 232 voxels, is visible across z = 22 to 42. Images are displayed on the MNI152 2-mm template in radiological orientation with a symmetric display range of ±5. (C) Post hoc ROI analysis of the right LC–left dlPFC cluster by age group. Mean LC–dlPFC connectivity did not differ reliably by age or group, whereas the inverse association between LC–dlPFC connectivity and positive bias was strongest in middle-aged adults. (D) Post hoc ROI analysis of the left LC–right angular gyrus/TPJ cluster by age group. Mean LC–angular gyrus/TPJ connectivity did not differ reliably by age or group,

A parallel post hoc analysis was conducted for the LC–angular gyrus/TPJ pathway **(Fig S5D)**. We extracted each participant’s mean gPPI connectivity estimate from the right angular gyrus/TPJ cluster identified in the left LC seed analysis and again examined both the mean connectivity estimate and its association with positive bias within each age group. As with the LC–dlPFC pathway, the mean LC–angular gyrus/TPJ connectivity estimate did not vary linearly with age, r = .07, p = .510, and did not show a reliable group effect, F(2, 87) = 0.80, permutation p = .454. The connectivity–bias association was negligible in children, r = .06, 95% CI [−.28, .34], p = .747, small and nonsignificant in emerging adults, r = −.21, 95% CI [−.49, .12], p = .286, and strongest in middle-aged adults, r = −.56, 95% CI [−.75, −.32], p = .001. This pattern suggests that the LC–angular gyrus/TPJ effect, like the LC–dlPFC effect, reflected an age-group difference in the functional relevance of LC-cortical connectivity to positive bias rather than an age-group difference in mean connectivity.

Because these ROI analyses were post hoc and the sample was cross-sectional, the age-group pattern should be interpreted descriptively. The results do not establish developmental change in LC-cortical coupling. Instead, they indicate that the relationship between LC-cortical connectivity and positive interpretation of emotional ambiguity was strongest in the middle-aged adult group and weaker or nonsignificant in the younger groups.

**Table S2.**
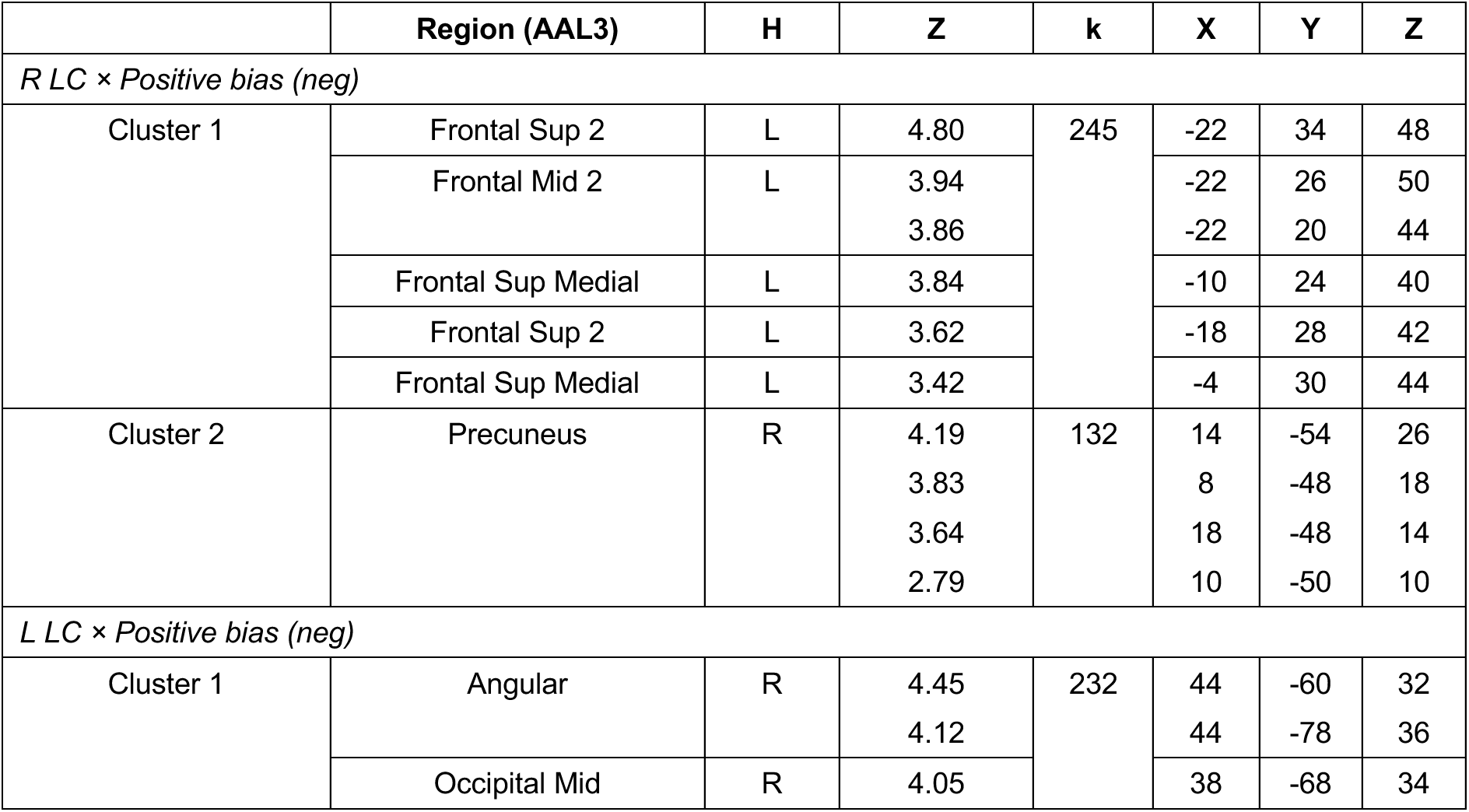
Brain regions showing LC connectivity associated with positive bias. Whole-brain generalized psychophysiological interaction, gPPI, analyses were performed separately for the right and left LC seeds. The regressor of interest was the interaction between LC seed time series and the individual-level positive-bias contrast, defined using PSE-anchored ambiguity weighting. Group-level inference used FSL FLAME 1+2 mixed-effects modeling, Z > 2.57, cluster-corrected p < .05. For each significant cluster, the peak voxel and up to five local maxima, separated by >8 mm, are listed. Regions were labeled using the Automated Anatomical Labeling atlas version 3, AAL3v1. In the main text, the left superior/middle frontal cluster, Cluster 1 from the right LC seed analysis, is referred to as left dlPFC based on its location in dorsal lateral prefrontal cortex. The right angular gyrus cluster, Cluster 1 from the left LC seed analysis, is referred to as right angular gyrus/TPJ based on its posterior inferior parietal location. H = hemisphere; Z = peak Z-statistic; k = cluster size in 2-mm isotropic voxels; x, y, z = MNI coordinates in mm.

### Trial-level DDM with Linear Ambiguity Weighting

As a sensitivity analysis for the Gaussian ambiguity weighting used in the main trial-level DDM, we refit the same model using a linear ambiguity-weighting function. In the main analysis, ambiguity was modeled as a Gaussian function centered on each participant’s PSE-nearest morph condition. Here, ambiguity declined linearly with ordinal distance from that participant-specific boundary: amb-linear(t) = max(0, 1 − |d(t)| / 3), where d(t) is the ordinal distance, in cope-selection-scale units, between the morph condition on trial t and the participant’s assigned PSE-nearest morph condition. For example, for a participant whose PSE-nearest condition was HA15, amblinear was 1.00 for HA15 trials, 0.67 for NE and HA35 trials, 0.33 for AN15 and HA60 trials, and 0 for more distant morph conditions. All baseline drift-diffusion parameters, drift slope, drift intercept, boundary separation, starting point, and nondecision time, were specified as in the Gaussian model. The same neural drift contributors were also retained: dlPFC, LC, and their interaction, along with their ambiguity-weighted terms. Thus, the only difference between the Gaussian and linear models was the shape of the ambiguity-weighting function applied to the ambiguity-moderated neural terms.

#### Group-level parameters

Group-level estimates were highly similar across the Gaussian and linear weighting schemes (**Table S3**). Baseline model parameters were essentially unchanged. Drift slope, drift intercept, boundary separation, and nondecision time were reliably different from zero in both models, whereas starting point was not.

For the neural drift contributors, no term reached conventional significance in either weighting scheme. The Gaussian model showed trend-level main-effect estimates for dlPFC, β_D_, p = .053, and LC, β_L_, p = .096. The linear model showed a similar pattern, with β_L_ at trend level, p = .074, and β_D_ not reliable, p = .120. The ambiguity-weighted neural terms were not reliable under either weighting scheme. Thus, the group-level parameter estimates did not indicate that the main findings depended on the specific Gaussian form of ambiguity weighting.

#### Individual-difference associations

Partial correlations between participant-level neural drift contributors and emotion-regulation measures, controlling for age, produced a pattern broadly consistent with the Gaussian model but with smaller effect sizes. The dlPFC ambiguity-weighted term, β_D·amb_,showed the strongest association with the ER composite among the neural contributors, although this effect was reduced to trend level under linear weighting, partial r | age = .19, p = .08. Reappraisal showed the same positive direction but was not reliable, partial r | age = .12, p = .26. Suppression was also not reliably associated with the dlPFC ambiguity-weighted term, partial r | age = −.14, p = .18. LC terms and LC × dlPFC interaction terms were uniformly weak and nonsignificant, consistent with the main Gaussian-weighted analysis; partial-correlation estimates are visualized in **Fig S6**

### Conclusion

The linear-weighting sensitivity analysis supported the same qualitative interpretation as the Gaussian-weighted trial-level DDM. The strongest individual-difference signal again appeared in the ambiguity-weighted dlPFC term, whereas LC terms and LC × dlPFC interaction terms remained weak or null. The dlPFC ambiguity-weighted association with the ER composite was smaller under linear weighting than under Gaussian weighting, partial r = .19, p = .08 versus Gaussian partial r = .23, p = .027. Thus, the trial-level dissociation between dlPFC and LC contributors was preserved, although the strength of the dlPFC ambiguity-moderated effect was somewhat sensitive to the form of ambiguity weighting.

**Fig S6.**
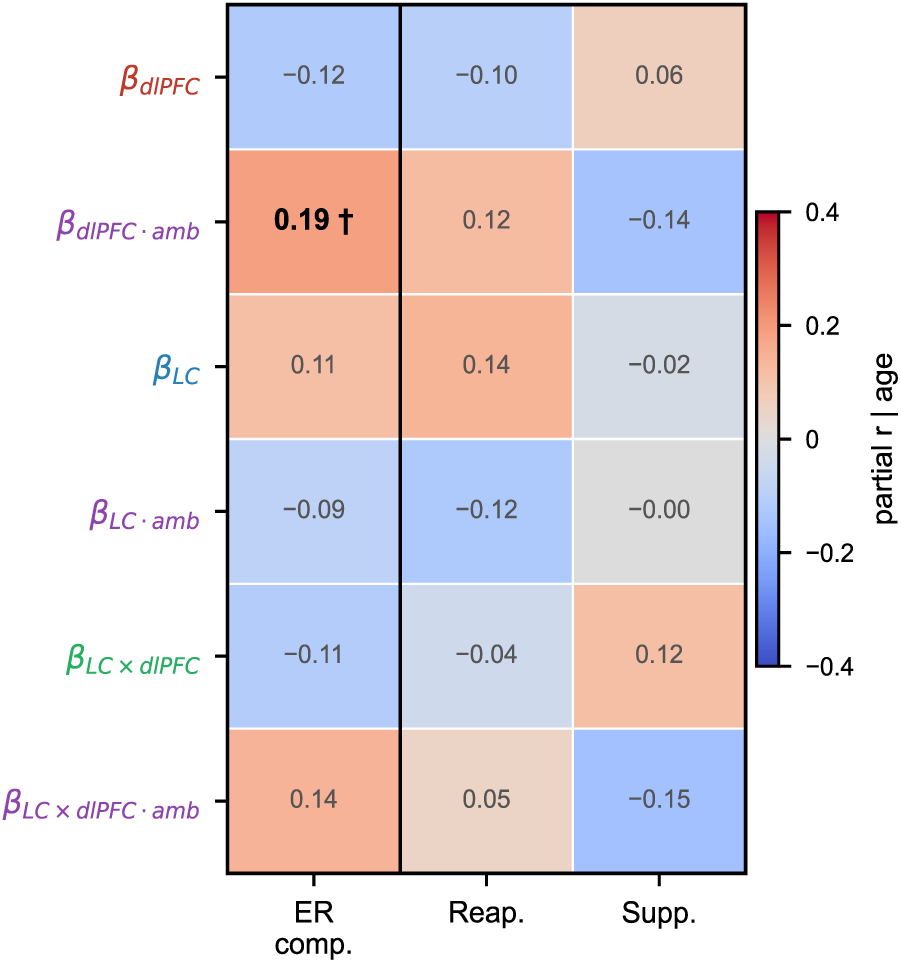
Partial-correlation heatmap from the linear-weighted trial-level DDM, controlling for age, N = 90. The heatmap relates the six neural drift contributors from the linear-weighted model to three emotion-regulation measures: ER composite, cognitive reappraisal, and expressive suppression. Row colors indicate the source term: red = dlPFC, blue = LC, green = LC × dlPFC interaction, and purple = ambiguity-moderated terms. The dlPFC ambiguity-weighted term, β_D·amb_, showed the strongest association with the ER composite, partial r | age = .19, p = .08, whereas LC terms were weak or null. The pattern was comparable to the Gaussian-weighted model reported in the main text, but with reduced effect sizes. †p < .10.

**Table S3.** Group-level trial-level DDM (M4) parameters under Gaussian and linear ambiguity weighting. Per-subject maximum-likelihood estimates from the trial-level drift-diffusion model (M4) were averaged across participants (N = 90) for the Gaussian weighting reported in the main text (amb(t) = exp(−d(t)^2 / 2)) and for the linear-weighting sensitivity fit (amb(t) = max(0, 1 − |d(t)| / 3)). Values are mean ± SE; t and p values reflect one-sample tests against zero. Baseline parameters (v_slope_, v_intercept_, B, x0, and nondecision time) index overall drift and diffusion dynamics independent of neural input. The six neural drift contributors comprise main effects and ambiguity-weighted slopes for standardized single-trial dlPFC BOLD (β_D,_ β_D·amb_), LC BOLD (β_L_, β_L·amb_), and their product (β_I_, β_I·amb_). At the group mean, the two weighting schemes yielded near-identical estimates across all parameters, and no neural contributor reached conventional significance in either fit.

| Parameter | Gaussian (main) | Linear (sensitivity) |
| --- | --- | --- |
| $V_{\text{slope}}$ | $3.922 \pm 0.112$, $p < .001$ | $3.959 \pm 0.111$, $p < .001$ |
| $V_{\text{intercept}}$ | $-0.335 \pm 0.060$, $p < .001$ | $-0.336 \pm 0.059$, $p < .001$ |
| B | $0.809 \pm 0.010$, $p < .001$ | $0.807 \pm 0.010$, $p < .001$ |
| $x_0$ | $0.010 \pm 0.011$, $p = .363$ | $0.008 \pm 0.011$, $p = .484$ |
| nondecision time | $0.290 \pm 0.008$, $p < .001$ | $0.293 \pm 0.008$, $p < .001$ |
| $\beta_D$ | $-0.041 \pm 0.021$, $p = .053$ | $-0.041 \pm 0.026$, $p = .120$ |
| $\beta_L$ | $0.030 \pm 0.018$, $p = .096$ | $0.039 \pm 0.021$, $p = .074$ |
| $\beta_I$ | $-0.021 \pm 0.024$, $p = .388$ | $-0.031 \pm 0.029$, $p = .294$ |
| $\beta_{D \cdot \text{amb}}$ | $-0.002 \pm 0.039$, $p = .959$ | $-0.000 \pm 0.048$, $p = .996$ |
| $\beta_{L \cdot \text{amb}}$ | $-0.015 \pm 0.037$, $p = .685$ | $-0.026 \pm 0.044$, $p = .552$ |
| $\beta_{I \cdot \text{amb}}$ | $-0.035 \pm 0.048$, $p = .464$ | $0.004 \pm 0.054$, $p = .945$ |

